# Multiplex genome editing eliminates the Warburg Effect without impacting growth rate in mammalian cells

**DOI:** 10.1101/2024.08.02.606284

**Authors:** Hooman Hefzi, Iván Martínez-Monge, Igor Marin de Mas, Nicholas Luke Cowie, Alejandro Gomez Toledo, Soo Min Noh, Karen Julie la Cour Karottki, Marianne Decker, Johnny Arnsdorf, Jose Manuel Camacho-Zaragoza, Stefan Kol, Sanne Schoffelen, Nuša Pristovšek, Anders Holmgaard Hansen, Antonio A. Miguez, Sara Petersen Bjorn, Karen Kathrine Brøndum, Elham Maria Javidi, Kristian Lund Jensen, Laura Stangl, Emanuel Kreidl, Thomas Beuchert Kallehauge, Daniel Ley, Patrice Ménard, Helle Munck Petersen, Zulfiya Sukhova, Anton Bauer, Emilio Casanova, Niall Barron, Johan Malmström, Lars K. Nielsen, Gyun Min Lee, Helene Faustrup Kildegaard, Bjørn G. Voldborg, Nathan E. Lewis

## Abstract

The Warburg effect is ubiquitous in proliferative mammalian cells, including cancer cells, but poses challenges for biopharmaceutical production, as lactate accumulation inhibits cell growth and protein production. Previous efforts to eliminate lactate production via knockout have failed in mammalian bioprocessing since lactate dehydrogenase has proven essential. However, here we eliminated the Warburg effect in Chinese hamster ovary (CHO) and HEK293 cells by simultaneously knocking out lactate dehydrogenase and regulators involved in a negative feedback loop that typically inhibits pyruvate conversion to acetyl-CoA. In contrast to long-standing assumptions about the role of aerobic glycolysis, Warburg-null cells maintain wildtype growth rate while producing negligible lactate. Further characterization of Warburg-null CHO cells showed a compensatory increase in oxygen consumption, a near total reliance on oxidative metabolism, and higher cell densities in fed-batch cell culture. These cells remained amenable for production of diverse biotherapeutic proteins, reaching industrially relevant titers and maintaining product glycosylation. Thus, the ability to eliminate the Warburg effect is an important development for biotherapeutic production and provides a tool for investigating a near-universal metabolic phenomenon.

## INTRODUCTION

Lactate is a key metabolite involved in many important processes. One of its prominent roles is in the Warburg effect^1^ in which highly proliferative cells (e.g., cancer, immune, and stem cells) exhibit high rates of glycolytic flux while secreting lactate even in the presence of oxygen^2–5^. Despite nearly a century of study, many questions remain surrounding its purpose, but it is believed to benefit cell proliferation^6^. However, this phenomenon of aerobic glycolysis has posed a considerable challenge in biotherapeutic protein production, since mammalian production cells often secrete high levels of lactate. Lactate accumulation is deleterious for cell growth, viability, protein production, and product quality, both directly from media acidification and indirectly through increased osmolarity as base is added to control culture pH^7–9^. Accordingly, a considerable amount of work for decades has aimed to control the Warburg effect in bioprocessing^10^.

In mammalian cells, lactate is produced by lactate dehydrogenase (Ldh). There are three Ldh isozymes: Ldha, Ldhb, and Ldhc, with different substrate specificities^10,11^, such as Ldhc’s preference for catalysis of the reverse reaction, forming pyruvate^12^. Ldha and Ldhb form homo- or heterotetramers with tissue specific distributions^13^. Ldha shows increased activity in many cancers^13,14^ and is present in the Chinese hamster ovary (CHO) Ldh complex^15^. While deficiencies in either LDHA^16–18^ or LDHB^19^ have appeared clinically in humans and are not lethal, efforts to delete Ldha in mammalian cell bioprocessing have been unsuccessful^20–22^, pointing to the essentiality of Ldh-mediated NAD^+^ regeneration. Many approaches have aimed to limit lactate accumulation in culture, including knockdown of Ldh^23,24^, replacement of glucose with alternate sugars^25–27^, controlled feeding strategies^7^, and others^10^; however none have successfully eliminated Ldh activity entirely.

Underpinning the enzymatic conversion of pyruvate to lactate by Ldh, the Warburg effect in mammalian cells is controlled by complex regulatory mechanisms^28^. Glucose is converted to pyruvate via glycolysis and pyruvate sits at the branch between fermentation to lactate by Ldh or oxidative metabolism starting with the pyruvate dehydrogenase complex (Pdh). A negative feedback loop involving several proteins and metabolites regulates the metabolic fate of pyruvate (Figure 1A). Specifically, the first step to oxidative metabolism, Pdh, can be regulated by four pyruvate dehydrogenase kinases (Pdks). Increases in the ratios of acetyl-CoA / CoA, ATP / ADP, or NADH / NAD^+^ will activate Pdk^29^. Pdks, when activated, phosphorylate and inhibit Pdh. Conversely, pyruvate inhibits Pdk1 and 2—thereby alleviating their inhibition of Pdh—but not Pdk3 or 4^29^. As excess flux through Pdh leads to Pdh inhibition, Ldh-mediated reduction of pyruvate to lactate functions as a release valve for the high glycolytic activity that occurs in proliferative cells. Thus, even in the presence of oxygen, glucose is not fully oxidized and lactate production is observed.

**Figure 1:**
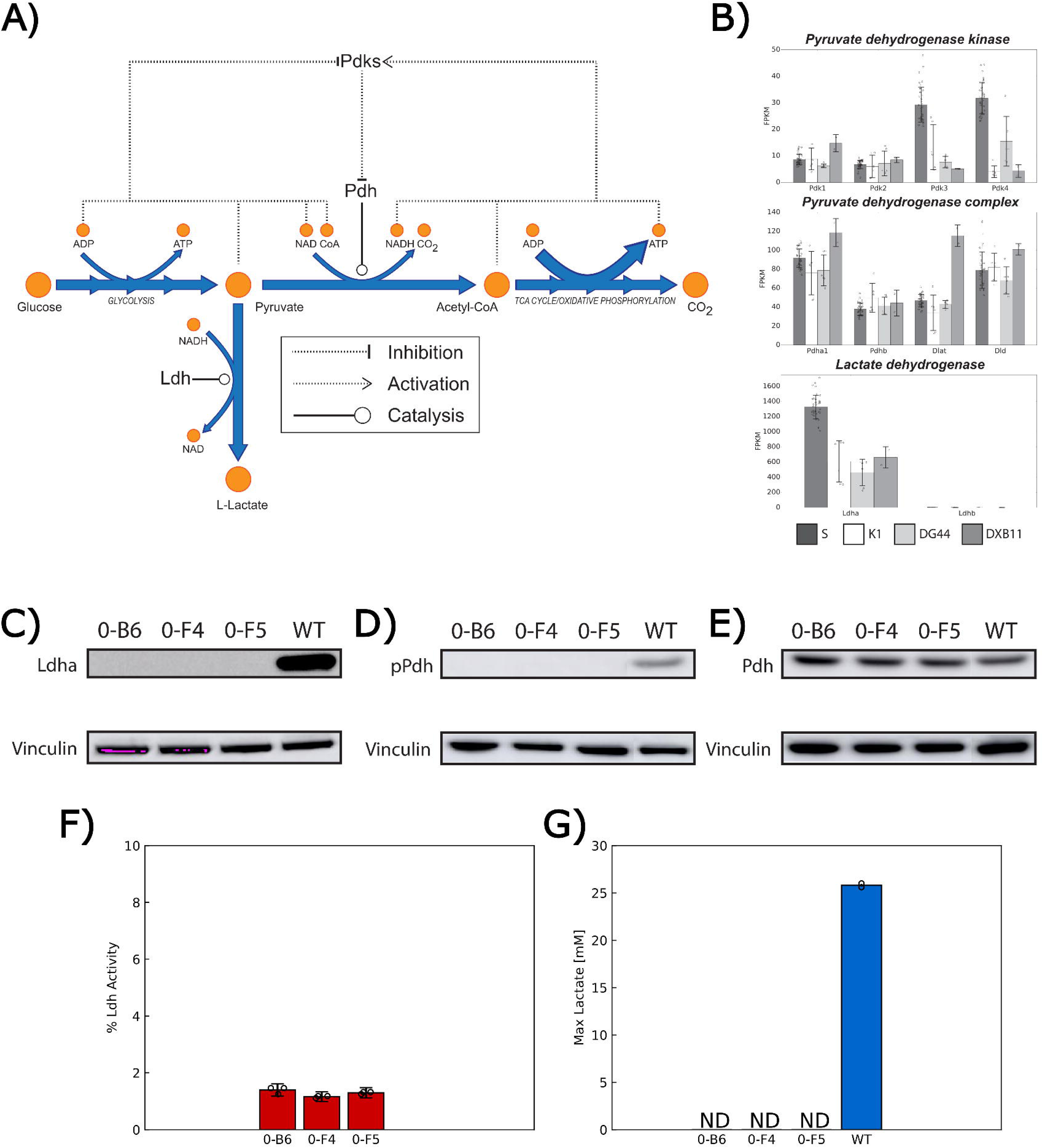
The Warburg effect is influenced by a regulatory circuit involving multiple metabolites and proteins and can be eliminated by multiplex knockout. (A) Pyruvate sits at a branch point between fermentation through lactate dehydrogenase (Ldh) or oxidative metabolism starting with the pyruvate dehydrogenase (Pdh) complex. Pyruvate dehydrogenase kinase isoforms (Pdks) are regulated by the products and substrates of the Pdh reaction, forming a negative feedback loop that reinforces an increase in lactate secretion when glycolytic flux is high. (B) Gene expression data from major CHO cell lines shows that Pdh complex subunits are all expressed, as are all four Pdk isoforms. Ldhb shows near zero expression (0 expression in CHO-S and DG44, below 0.1 FPKM in CHO-K1 and DXB11) while Ldha expression is high. All gene expression values are given in fragments per kilobase per million reads (FPKM), data sources are provided in Supplementary Table 4. (C) Simultaneously targeting Ldha and the Pdk genes for knockout using CRISPR/Cas9 leads to no detectable Ldha expression, as verified by chemiluminescent Western blot. (D) No phosphorylation of Pdh at S293 is detected in knockout lines. (E) Total Pdh remains stable in knockout lines. (F) Enzymatic assay verifies a loss of lactate dehydrogenase activity in knockout lines (shown here as percent of wildtype Ldh activity, n=3 technical replicates per cell line). (G) Measurement of maximum lactate levels during growth in batch culture (Figure 2, clones with 4 Pdk knockouts) shows that lactate cannot be detected (n=2 shake flasks per cell line). Data shown as mean ± standard deviation (uncertainty due to variance in background in (F) accounted for via propagation of error). Gene abbreviations are as follows: Pdk1-pyruvate dehydrogenase kinase 1, Pdk2-pyruvate dehydrogenase kinase 2, Pdk3-pyruvate dehydrogenase kinase 3, Pdk4-pyruvate dehydrogenase kinase 4, Pdha1-pyruvate dehydrogenase E1 alpha 1 subunit, Pdhb-pyruvate dehydrogenase E1 beta subunit, Dlat-dihydrolipoamide S-acetyltransferase, Dld-dihydrolipoamide dehydrogenase, Ldha-lactate dehydrogenase A, Ldhb-lactate dehydrogenase B.

Here we report that removing the feedback loop around pyruvate permitted the complete elimination of Ldh activity in mammalian cells without negatively impacting growth. We achieved this through the homozygous knockout of lactate dehydrogenase(s) and the four Pdks in CHO and HEK293 cells. The cells remain proliferative while consuming significantly less glucose. We further characterize the phenotypic response of eliminating the Warburg effect and demonstrate the cells remain amenable for use in the traditional cell line generation process of the biopharmaceutical industry. Additionally, we show that introduction of the Warburg-null phenotype into monoclonal antibody (mAb) producing cell lines maintains product titers and important quality attributes such as glycosylation. These results suggest that the Warburg effect is not essential to obtain biomass precursors to fuel mammalian cell growth and protein production in bioprocessing, and the removal of the effect opens new opportunities for improving production of biotherapeutics.

## RESULTS

### Ldha can be knocked out when concurrently targeting Pdk1, 2, 3, and 4

Across 54 CHO RNA-Seq samples from 4 CHO cell lineages, all four Pdk isozymes and core Pdh subunits are expressed. While Ldha is highly expressed, Ldhb is silenced, indicating that the Ldh enzyme is made exclusively of Ldha subunits (Figure 1B). Previous efforts to knockout Ldha in CHO cells have been unsuccessful^20–22^, possibly because the knockout of Ldha would lead to accumulation of pyruvate, which cannot be fully processed by Pdh due to activation of Pdks inhibiting Pdh. We thus aimed to remove Ldha and the Pdk-mediated feedback loop by simultaneously removing Ldha and all four Pdks. We hypothesized that this would relieve Pdk-mediated inhibition of Pdh and allow uninhibited flux of pyruvate through Pdh. Therefore, using CRISPR/Cas9, we simultaneously targeted all five genes in the CHO-S (WT) cell line (Supplementary Table 1). Consistent with our hypothesis, we isolated several clones with frameshift mutations in all alleles of the five genes (Supplementary Tables 2-3).

Further characterization of the five gene knockout clones confirmed that the Warburg effect was eliminated. We measured Ldha by Western blot, showing that protein expression was undetectable (Figure 1C). We also assessed Pdk activity by measuring phosphorylated Pdh1a levels via Western blot and mass spectrometry. As expected, no detectable phosphorylation of Pdh1a was observed; additionally, we note total Pdh1a increased slightly (Figure 1D/E, Supplementary Figure 2). We further tested the cell lysates from knockout clones for Ldh activity and found that it was reduced to near zero (Figure 1F). Finally, lactate was undetectable when assayed by the industry-standard BioProfile 400 (Figure 1G). We further employed more sensitive methods to quantify the decrease in lactate levels in a variety of clones and culture conditions. We found that maximum lactate concentrations obtained were in the sub-millimolar range for clones 0-F4 and 0-F5, while clone 0-B6 reached single millimolar concentrations (Supplementary Figure 1). When compared to values observed in wildtype or Warburg-positive clones/pools (maximum concentrations ranging between ∼13-135 mM lactate) it is apparent that the deletion of Ldha effectively abolished Warburg metabolism in these cells.

### Knocking out subsets of Pdks permits Ldha knockout in CHO and HEK 293

In addition to clones with all five genes knocked out, we found a minority of clones with frameshift mutations in Ldha, while retaining some wildtype Pdk alleles. This was surprising, as previous work targeting Pdk1-3 with siRNAs could not isolate a complete Ldha knockout^21^. Therefore, we conducted four more rounds of CRISPR/Cas9-mediated engineering, targeting all combinations of three Pdks alongside Ldha (e.g., Ldha + Pdk 1, 2, and 4) to verify that the Warburg effect could be eliminated without knocking out all Pdks. In total we single-cell sorted 378 clones (93-95 clones per 3-Pdk combination) for genotyping by next-generation sequencing.

Between these clones and those obtained from targeting all Pdks and Ldha simultaneously (515 clones in total), we found diverse clones with different combinations of gene knockouts. First, consistent with previous work^20–22^, we could not obtain stable, viable clones with only homozygous Ldha frameshift mutations. Second, we found several clones harboring complete Ldha knockouts alongside diverse combinations of Pdk knockouts, including some with only 1, 2, or 3 Pdks eliminated (Supplementary Tables 2-3), and separately verified that the knockout of Pdk2 was sufficient to allow subsequent knockout of Ldha and eliminate Warburg metabolism in CHO cells (data not shown).

Finally, we attempted the original knockout strategy in a suspension, serum-free adapted CHO-K1 cell line and a HEK293 cell line, the latter of which required additional targeting of LDHB. This resulted in clones with some or all Pdks knocked out alongside Ldh(s) (Supplementary Tables 5, 6, 8, 9), suggesting that this engineering strategy is a generalizable way to eliminate the Warburg effect.

Further characterization of these clones confirmed knockout of Ldh(s) and elimination of the Warburg effect (Supplementary Figure 2), leading to growth rates comparable to wildtype with substantially reduced glucose uptake (Figure 2A/B, Supplementary Figures 3/4). Interestingly, we saw variation in the pattern of Pdh expression and phosphorylation: While most clones showed a decrease in their fraction of phosphorylated Pdh and increase in total Pdh (Supplementary Figure 5), this was not universally true. Deeper characterization of these clones is necessary to determine how altered levels of phosphorylation vs. total Pdh levels interact to permit the Warburg-null phenotype to emerge in incomplete Pdk knockouts.

**Figure 2:**
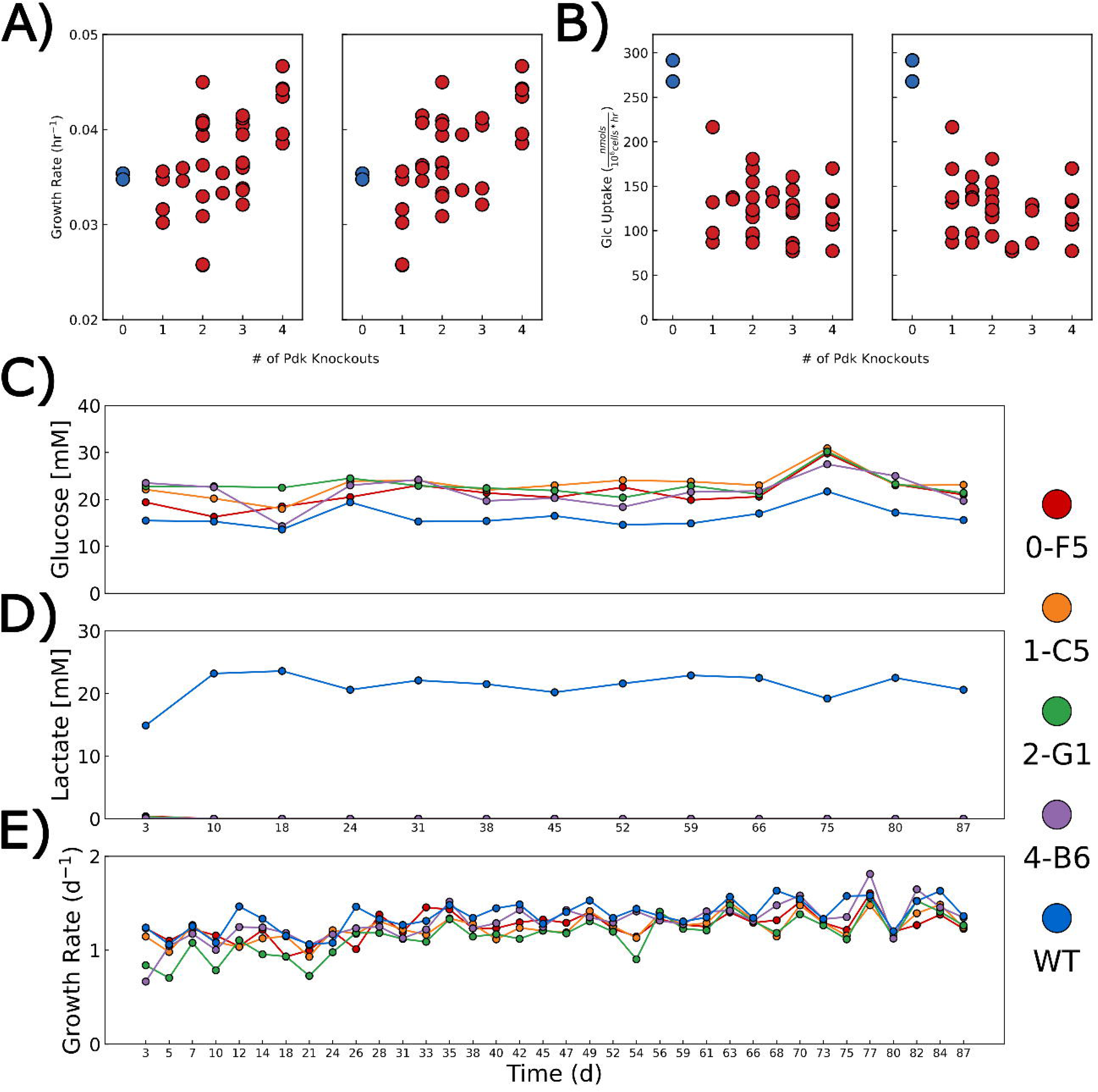
Growth rate, glucose uptake, and long-term passaging of Warburg-null clones. Warburg-null cell lines (n=2 shake flasks for each clone) remain proliferative with (A) a growth rate comparable to WT while (B) drastically reducing their glucose uptake rate. This effect is observed whether in-frame mutations are counted as knockouts (left) or wildtype (right). The growth of wildtype CHO-S is shown in blue. Long-term passaging over approximately 87 days shows that Warburg-null clones (C) have higher residual glucose in culture while (D) not producing lactate and (E) maintain a comparable growth rate to WT cells (n=1 shake flask for each cell line in long-term passaging experiment).

### Eliminating the Warburg effect rewires central metabolism without affecting growth rate

It has been hypothesized that the Warburg effect is necessary for cell proliferation by allowing production of biomass precursors^6^. While our initial test did not show an impact on growth rate in any Warburg-null clones, to ensure that the Warburg-null phenotype would not be selected against over time (if there is a growth benefit to lactogenic behavior), we subjected the 0-F5 clone and clones with Ldha and only 1, 2, or 3 Pdks knocked out (4-B6, 2-G1, and 1-C5, respectively) to long-term routine passaging. After 87 days of passaging, all non-WT cells remained stably Warburg-null (Figure 2C-E). Thus, the Warburg effect is dispensable for cell proliferation.

Another hypothesized purpose of the Warburg effect is to increase ATP generation through substrate-level phosphorylation from high glycolytic flux^6^. Thus, we investigated if the cells need aerobic glycolysis to meet their ATP needs, and if there is a compensatory response to maintain ATP production to offset the decreased glycolytic flux in Warburg-null cells. To evaluate this, 0-F5 and WT cells were grown in a controlled bioreactor for deeper characterization (Figure 3A). Again, the Warburg-null phenotype did not show any effect on growth rate, demonstrating that the ATP production from high glycolytic flux is not essential for normal growth. However, owing to the vast difference in ATP yield between oxidative metabolism and substrate level phosphorylation from glycolysis, large decreases in glycolytic flux could be balanced by small increases in oxygen consumption, if the mitochondria could support increased oxidative metabolism. Accordingly, we observed a slight increase in the oxygen uptake rate for Warburg-null lines (Figure 3B), which we confirmed using the Seahorse Analyzer (Figure 3C). The latter also revealed a lack of compensatory increased glycolysis upon treatment with electron transport chain inhibitors, indicating near complete reliance on mitochondrial metabolism for energy. Indeed, Warburg-null cells were more rapidly killed by BAM15 (a strong mitochondrial uncoupler^30^) than wildtype cells (Figure 3D). To confirm increased oxidative metabolism, in separate experiments (Figure 3E, Supplementary Figure 7A), ^13^C labeled glucose was fed in mid-exponential phase and labeling measured over time via LC-MS (Figure 3G) or GC-MS (Supplementary Figure 7B) where we observed increased labeling of TCA cycle metabolites with both methods. Finally, in a 4th experiment (Figure 3F), we observed an increase in the intracellular pyruvate and TCA cycle intermediate levels (Figure 3G). These results are all consistent with increased entry of glucose-derived pyruvate into the mitochondria for full oxidation. Thus, while Warburg metabolism does not appear to be essential, Warburg-null cells display a compensatory increase in--and a reliance on--mitochondrial oxidative metabolism for ATP generation.

**Figure 3:**
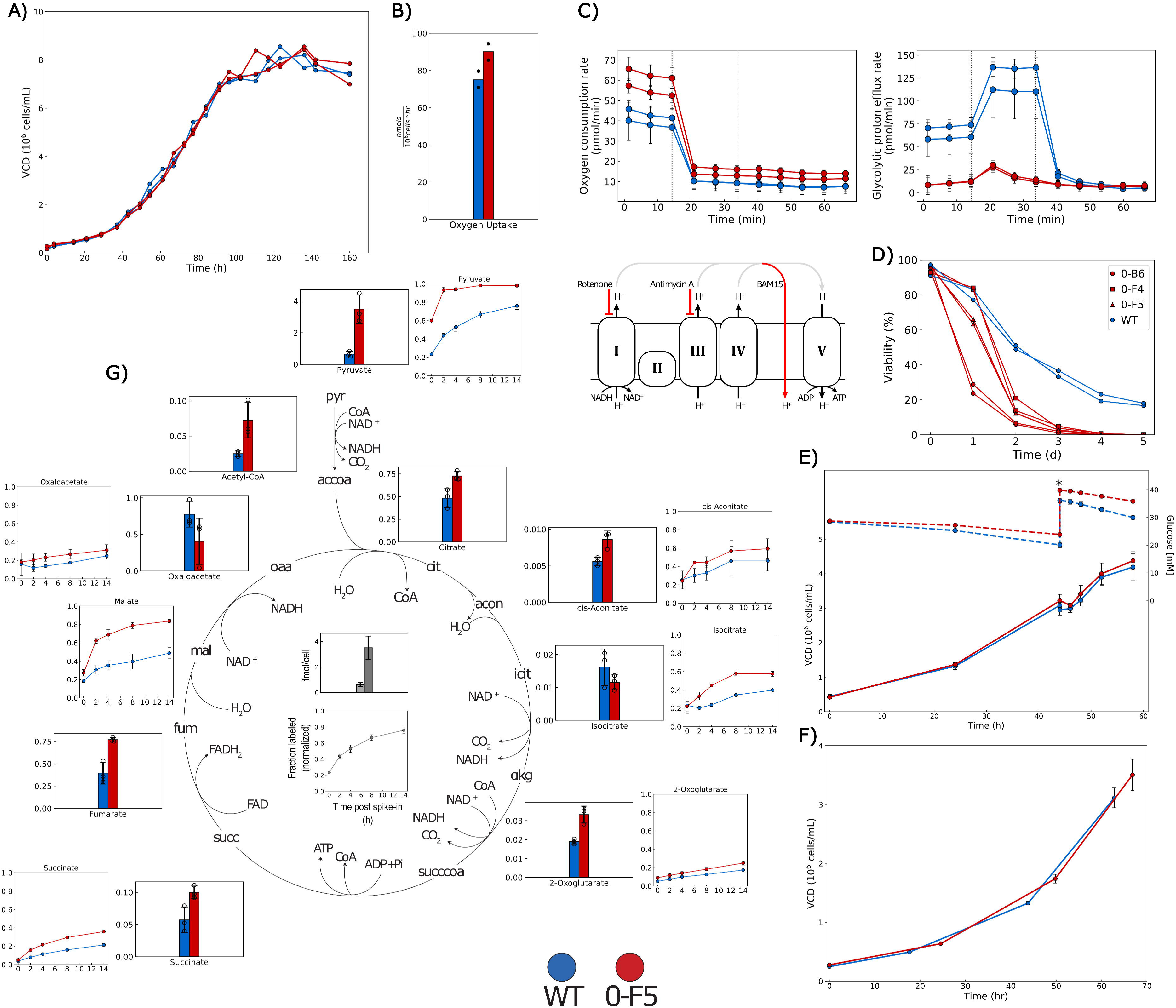
Warburg-null cells increase oxidative metabolism. (A) WT and 0-F5 (n=2 bioreactors per cell line) cells grown in pH and DO controlled bioreactors in batch culture showed near identical growth profiles. (B) Warburg-null cells from (A) showed increased oxygen uptake (calculated between t=20 and 68 hrs, see Methods). (C) Cells (2 independent vial thaws per cell line, 23 wells per vial) were assayed using the Agilent Seahorse Glycolytic Rate Assay to calculate oxygen consumption rate (left) and glycolytic proton efflux rate (right), an estimate of the glycolytic contribution to extracellular acidification. Warburg-null cells show increased oxygen consumption and little glycolytic proton efflux. The first dotted line (∼14 min) denotes the addition of rotenone and antimycin A while the second (∼33 min) denotes the addition of 2-deoxyglucose. Warburg-null cells do not compensate for the cessation of oxygen consumption with an increase in glycolytic activity. (D) Warburg-null cells show complete reliance on oxidative metabolism as treatment with BAM15 (a mitochondrial uncoupler^30^) rapidly kills Warburg-null cells while some wildtype cells survive (n=2 wells per cell line). A schematic of the electron transport chain (ETC) and the mechanism of action of rotenone, antimycin A, and BAM15 are depicted. I-V denote the different protein complexes of the ETC. (E) Cells were grown in bioreactors (n=4 bioreactors per cell line) and when cells reached ∼3×10^6^ cells/mL, ^13^C labeled glucose was spiked in (timepoint indicated by *). Cultures were then sampled 5 times over the following 14 hours. (F) Cells were grown in bioreactors (n=3 bioreactors per cell line) and harvested at the final time point shown for transcriptomic and endometabolomic analysis. (G) Intracellular levels for TCA cycle metabolites are markedly increased overall (bar charts). Labeling of TCA cycle metabolites following the addition of ^13^C-labeled glucose also increased (line graphs). Abbreviations-pyr: pyruvate, accoa: acetyl-CoA, cit: citrate, acon: aconitate, icit: isocitrate, LJkg: alpha-ketoglutarate/2-oxoglutarate, succcoa: succinyl-CoA, succ: succinate, fum: fumarate, mal: malate, oaa: oxaloacetate.

### Cells show dysregulated redox metabolism but diverse transcriptional responses to elimination of the Warburg effect

Since Ldha regenerates NAD+, we investigated how its knockout impacts the redox balance of the cell. Intracellular metabolomics measured during mid-exponential growth showed a dramatic decrease in the NAD/NADH ratio (Figure 4A) stemming from a large increase in NADH levels that was not fully compensated for by increased NAD+ levels (Figure 4B), consistent with the loss of Ldha-mediated NADH recycling. Possibly to alleviate this decrease, the 0-F5 cell line showed a substantial increase in the intracellular glycerol-3-phosphate pool (Figure 4C) and upregulation of *Gpd1*, an alternative NAD+ regenerating enzyme (Figure 4D). Transcriptomic data from other clones (0-B6 and 0-F4) and from the CHO-K1-derived Warburg-null line also suggests dysregulation of NAD/NADH levels, with all clones showing altered expression levels of several metabolic enzymes utilizing NAD/NADH and/or metabolic precursors and evidence of *Sirt1* activation (Figure 4D).

**Figure 4:**
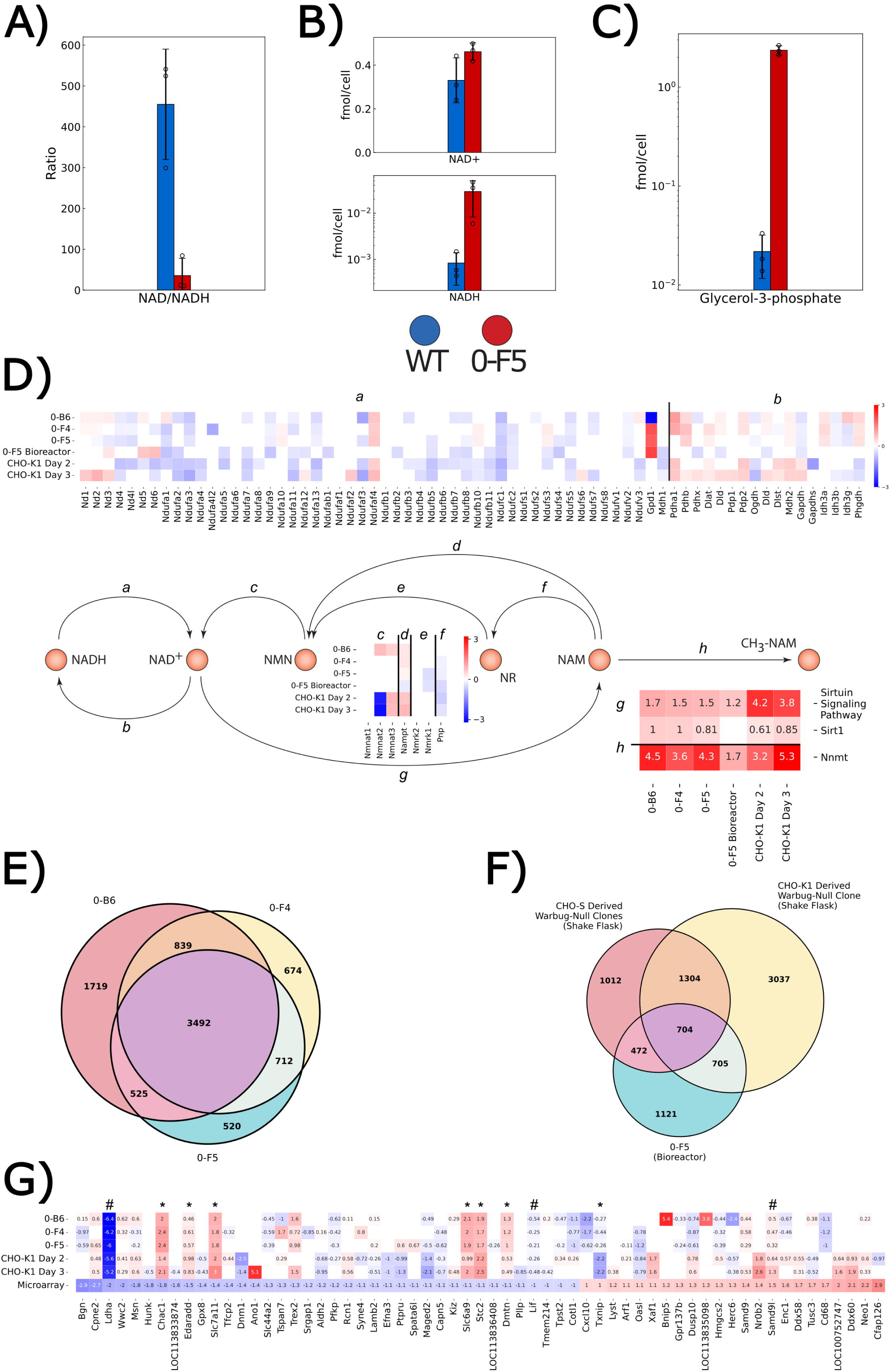
Warburg-null cells show perturbations in redox state and metabolism, though these changes may be clone, cell line, and organism specific. (A) Loss of Ldha considerably decreases the NAD/NADH ratio in Warburg-null cells due to (B) a large increase in NADH levels and a minor increase in NAD+ levels. This is accompanied by (C) intracellular accumulation of glycerol-3-phosphate compared to WT cells. (D) Across cell lines and conditions, Warburg-null cells show perturbations in genes involved in redox metabolism. Genes are labeled based on their function in the metabolic map: a/b utilize NAD/NADH as cofactors, c/d/e/f consume or produce NAD and its biosynthetic precursors, g consumes NAD, and h irreversibly removes NAM from the NAD precursor pool. All data show the log_2 f_old change of the Warburg-null line compared to the wildtype, except for the Sirtuin Signaling Pathway (z-score from Ingenuity Pathway Analysis). (E) Each Warburg-null clone grown in batch culture shows clone-specific differentially expressed genes (DEGs). (F) DEGs shared in all 3 clones from (A) are not consistently differentially expressed when a clone is grown in different culture conditions (0-F5 bioreactor) or when the clone is derived from a different CHO cell line (CHO-K1 DEGs are the intersection of DEGs from days 2 and day 3). (G) DEGs differ even more when compared to a Warburg-null human cancer cell line (LS174T)^31^ assayed via microarray (values reported are log_2_ fold changes; Benjamini-Hochberg FDR < 0.05). # marks genes that are differentially expressed in the same direction as the human cancer cell line in all CHO samples. * marks genes that are differentially expressed in all CHO samples but in the opposite direction as seen in LS174T cells. Note: Samples in (A), (B), (C), and the 0-F5 Bioreactor experiment in (D) are from the final timepoint in Figure 3F (n=3 bioreactors per cell line). Samples 0-B6, 0-F4, 0-F5 are from day 3 of cultures in Supplementary Figure 8 (n=3 shake flasks per cell line). CHO-K1 Day 2 and CHO-K1 Day 3 are from samples taken at day 2 and day 3, respectively, from the experiment depicted in Supplementary Figure 3 (n=3 shake flasks per cell line). All comparisons were of the Warburg-null clone(s) against paired Warburg-positive samples, DEG cutoff for all comparisons: Benjamini-Hochberg FDR < 0.05. Abbreviations-NMN: Nicotinamide D-ribonucleotide, NR: N-ribosyl-nicotinamide, NAM: nicotinamide, CH_3_-NAM: N1-methyl-nicotinamide.

Despite all tested clones and conditions showing perturbations around redox metabolism, a common, conserved transcriptional response to elimination of the Warburg effect did not appear to be present. *Gpd1*, for example, shows increased expression in 2 of 3 CHO-S derived clones but decreased expression in the 3rd. Furthermore, no change in *Gpd1* is seen in the CHO-K1 derived Warburg-null clone (Figure 4D), where *Gpd1* shows minimal expression in both wildtype and knockout lines. Even among clones in identical culture conditions there were dramatic differences in transcriptional profiles, with a large number of unique differentially expressed genes in clone 0-B6 (Figure 4E). This difference was also observed for different growth media/conditions and especially apparent when compared to the differentially expressed genes in the CHO-K1 derived Warburg-null clone (Figure 4F). Finally, we compared the transcriptional changes observed in our clones to a study that generated a Warburg-null LS174T colon cancer line via simultaneous knockout of LDHA and LDHB^31,32^, resulting in reduced growth. Out of 192 genes showing strong differential expression (|log_2_ fold change|>1) in LS174T, we found clear homologs for 147 genes in CHO; however only ∼40% showed statistically significant differential expression in *any* CHO cell line and only 3 were differentially expressed in the same direction as the human cancer cell line in all CHO samples (one of these genes being Ldha), while 7 genes showed the opposite pattern (i.e., differentially expressed in all CHO samples but in the opposite direction seen in LS174T cells (Figure 4G). Together, these findings strongly suggest that there may be several cellular states that permit a Warburg-null phenotype to arise in mammalian cells.

### Warburg-null cells are compatible with industry-standard cell line generation protocols

The elimination of the Warburg effect addresses a fundamental challenge in mammalian bioprocessing. We observed no negative effects on growth rate in batch culture (Supplementary Figures 5,6,8) and a prolonged growth period and higher maximum VCD in fed-batch, consistent with a lack of osmolite-induced entry to stationary phase (Supplementary Figure 9). Since these cells would ultimately be used for biomanufacturing, we investigated how the Warburg-null phenotype impacts the process of cell line development (CLD).

Industrial cell line development procedures involve transgene integration, selection, and amplification using selectable markers. These result in polyclonal pools producing the transgene, from which stable clones are identified and used for production^33^. Ideally, engineered host cell lines would be compatible with existing CLD workflows, but previous work showed that shRNA-mediated downregulation of Ldha resulted in cells unable to survive the established selection and amplification processes^33,34^. These results raise concerns that the knockout of Ldha and Pdk-mediated Pdh regulation may result in a similarly unsuitable host cell line. Fortunately, when we subjected our Warburg-null clones to an identical selection and amplification process, we successfully generated polyclonal pools of mAb-producing cells that behaved similarly to WT cell-derived pools (Supplementary Figure 11). Thus, Warburg-null cells are compatible with established gene amplification protocols.

### Warburg-null cells can produce diverse therapeutic proteins

To verify that our knockout of Ldha and the Pdk genes did not disrupt the protein secretion capacity of the cells, we transfected the Warburg-null clones with diverse biotherapeutics including Enbrel as well as non-Fc domain containing drugs, including erythropoietin, C1 esterase inhibitor, and growth/differentiation factor 5. We successfully obtained clonal cell lines producing each protein (Supplementary Figure 13), indicating broad applicability of this phenotype for protein production. Finally, to verify the ability to obtain clones with meaningful product titers, we used BESTCell^TM^ technology^35^ (The Antibody Lab, Vienna, Austria) to generate Herceptin and Rituximab producing clones and obtained titers >1 g/L after a 10 day fed-batch screening process in deepwell plates (Supplementary Figure 14). Two Warburg-null Rituximab-producing clones were selected for further evaluation and found to be compatible with a prototypical high-intensity fed-batch process in bioreactors, reaching industrial titer levels (up to ∼3 g/L) and maintaining viability for two weeks—without requiring base addition to control pH as a result of the lack of lactate production (Figure 5A-C, Supplementary Figure 15). Thus, Warburg-null cells are compatible with established cell line development protocols and show suitability as host cell lines.

**Figure 5:**
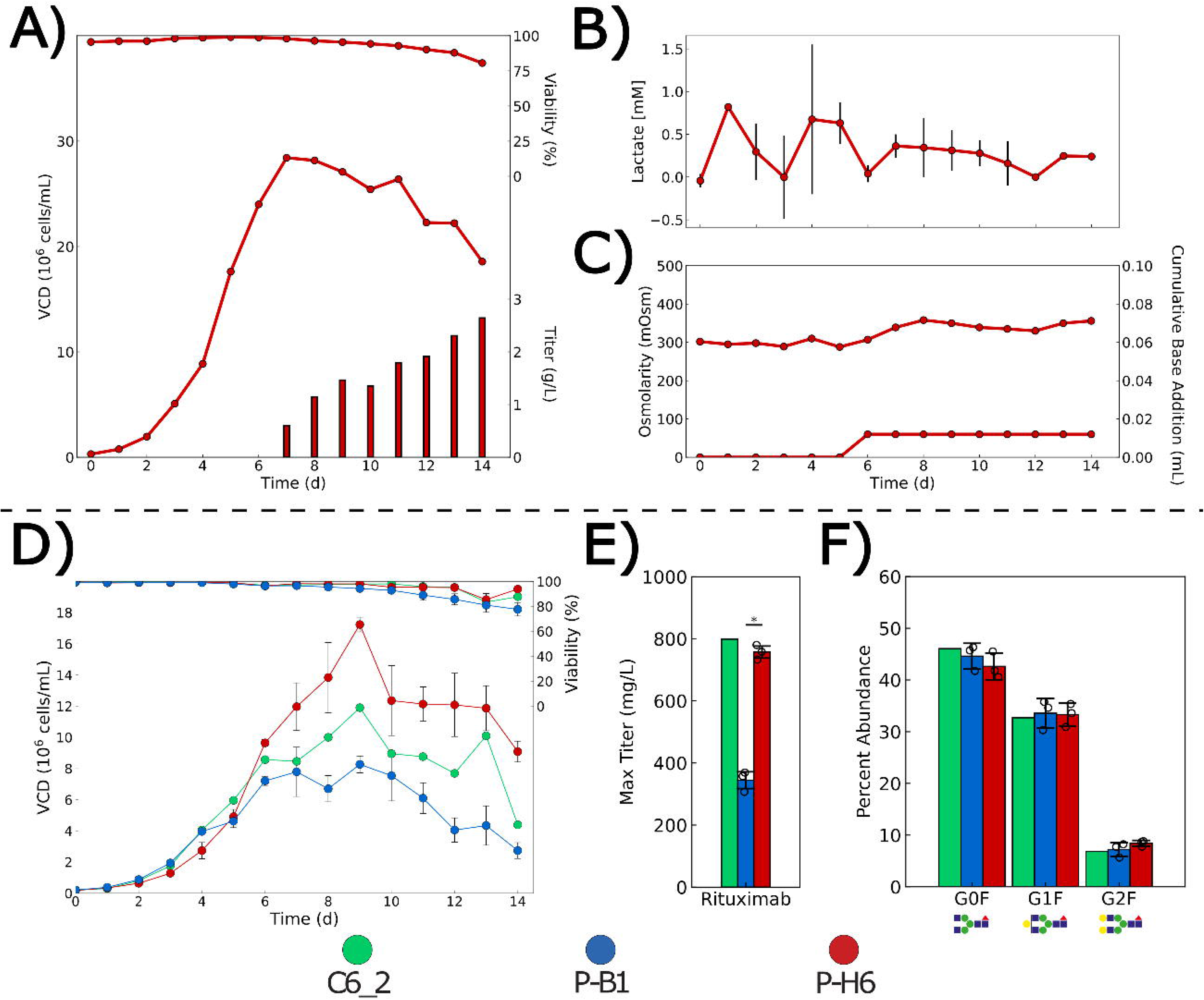
Warburg-null cells can be used as a new host cell line or the phenotype can be introduced into existing production clones. (A-C) The best performing individual vessel from a fed-batch evaluation in ambr15 bioreactors of the top 2 clones generated from the 0-F5 Warburg-null clone (see Supplementary Figure 15 for all vessels). (A) Growth, viability, and rituximab titer. (B) Lactate, (C) osmolarity, and base addition over the course of culture. Maximum vessel volume was 15 mL. Due to the lack of base addition, culture osmolarity varies minimally. (D-F) Separately, parental (C6_2, green, n=1 bioreactor), mock control (P-B1, blue, n=3 bioreactors), and Warburg-null (P-H6, red, n=3 bioreactors) cells producing rituximab were grown in fed-batch culture. (D) Knockout cells showed a prolonged period of exponential growth compared to both the parental and control line (see Supplementary Figure 1A for maximum lactate concentrations for all clones). (E) Knockout cells were able to maintain (vs. parental) or improve (vs. control) product titer (* indicates p<0.05 as determined by a two-sample two-tailed t-test between mock and Warburg-null clones). (F) Warburg-null clones maintained comparable glycosylation of rituximab at day 14 of the culture. (* indicates significance at a Benjamini–Hochberg FDR < 0.05, following a two-sample two-tailed Welch’s t-test for mock vs. Warburg-null comparisons). Data shown as mean ± standard deviation.

### The Warburg-null phenotype can be introduced into existing mAb-producing lines

Generating high productivity cell lines is time-consuming, so it would be advantageous to introduce the Warburg-null phenotype into existing drug-producing CHO cell lines with undesirable lactate profiles (prior to generation of material for clinical trials). To demonstrate this, we targeted all Pdks and Ldha in a Rituximab-producing CHO cell line (see Supplementary Methods) and isolated two clones with Ldha—albeit not all Pdks—knocked-out (Supplementary Tables 2-3). In parallel, as controls for the cell engineering process, we isolated five clones (referred to here as “mock”) derived from transfection with only Cas9.

Following initial characterization and media screening (Supplementary Figure 16), we found that--consistent with the non-producing Warburg-null clones (Supplementary Figure 9)--the mAb-producing cells without Ldha exhibited improved growth behavior compared to both the parental and mock lines (Figure 5D). Protein titer was maintained at similar levels to the parental line (and increased compared to the control clone, Figure 5E). Furthermore, product glycosylation was maintained (Figure 5F, Supplementary Figure 17). Thus, the Warburg-null phenotype can be introduced into existing biopharmaceutical cells without losing protein production or product quality.

## DISCUSSION

Here we report that the Warburg effect can be eliminated via removal of lactate dehydrogenase and peripheral regulators in a proliferative mammalian cell line. The engineered CHO cells drastically decreased their glucose consumption alongside the near-zero lactate secretion. Surprisingly, the cells remain rapidly proliferative, and, when grown in fed-batch, exhibit an extended growth phase. This likely results from avoiding medium hyperosmolarity, since base addition is not needed to neutralize the lactic acid.

As CHO cells are the primary workhorse of the biopharmaceutical industry, advantageous phenotypes must be amenable to existing workflows for generating new protein producing cell lines or be introducible into existing lines without deleterious effects. We demonstrated that the Warburg-null phenotype satisfies both criteria. As a host cell line, polyclonal pools producing recombinant antibody were generated following standard industrial workflows. These pools behaved similarly to pools derived from wildtype CHO-S cells, in contrast to previous work showing Ldha-downregulated cells could not survive the standard workflow^34^. Furthermore, clones derived from pools producing other biotherapeutic proteins also behaved similarly to their WT-derived counterparts. Ultimately, industrially-relevant product titers were obtained using Warburg-null clones as the starting cell line. Introducing the phenotype into a biopharmaceutical-producing CHO cell line improved growth by prolonging the exponential phase and maintained or improved product titers while retaining appropriately galactosylated glycans. We note that these improvements were seen in processes designed around Warburg positive cells (e.g. glucose-rich media where the anticipated entry into stationary phase is osmolarity linked). Thus, we anticipate that further bioprocess optimization around the Warburg-null phenotype would further enhance these benefits. For example, without lactate secretion and the associated base addition to control pH, culture osmolarity is less of a constraint, enabling bioreactor runs with a longer growth phase^7^ or overall duration with an appropriately tailored bioprocess. There will be additional opportunity for optimizing media formulation and adjusting feeding of amino acids, owing to the interplay between the metabolism of these compounds. For example, excess amino acids are often added, since they reduce aerobic glycolysis^36^. However amino acid catabolism generates ammonia and other growth-inhibitory compounds^37^. Without lactate secretion, amino acids could be maintained at lower concentrations and used to primarily feed protein synthesis^37^. These potential advantages can be explored through further studies to test the long-term implications of the removal of the Warburg effect.

The application of our approach to diverse mammalian cells (CHO and HEK) highlights its utility as a novel model system in which others can address fundamental questions regarding the Warburg effect. That is, the ubiquity of aerobic lactate production in proliferative eukaryotic cells has led to a rich field of study to understand how and why this phenomenon occurs^6,38^. However, while numerous studies have inhibited lactate production to varying degrees^14,15,21,39–41^ (e.g. using siRNA, chemical inhibitors, or CRISPR), the lack of proliferative Warburg-null cell lines has hindered full exploration of this phenotype. While we focused primarily on the biotechnological relevance of the Warburg effect, our study provides interesting insights into outstanding hypotheses and debates in the broader scientific discipline. First, it is commonly assumed that immortalized cells such as CHO and cancer have either dysfunctional mitochondria or insufficient mitochondrial capacity for needed ATP generation and therefore the Warburg effect is needed to support the energetic demands of growth, cell maintenance, and other activities^6,38,42^. However, while the Ldha knockout resulted in decreased glucose uptake and glycolytic flux, the cells channel more glucose into oxidative metabolism. Indeed, we observed an increase in oxygen uptake and increased labeling of TCA cycle metabolites after feeding with ^13^C labeled glucose. Thus, the Warburg effect is not necessarily a response to mitochondrial insufficiency^43^, and Warburg-null cells can compensate for the loss of ATP generated via high glycolytic flux in wildtype cells. Second, many studies have suggested that increased glycolytic metabolism is necessary for producing biomass precursors for proliferation seen in cancer, T-cell activation, and embryonic development^6,44,45^. However, we found that elimination of the Warburg effect does not appear to harm the cells or hinder their ability to proliferate. While we can therefore confidently say that aerobic lactate production is not needed to meet energetic or biosynthetic demands of proliferative cells, the evolutionary benefit to this ubiquitous phenotype remains ripe for investigation. The genome-editing strategy proposed here provides an approach to develop new cell models to explore such questions.

In conclusion, we found the elimination of the Warburg effect is feasible in mammalian cells by eliminating lactate dehydrogenase activity. While prior studies showed that Ldha in CHO cells was essential^20–22^, we found that its essentiality in CHO cells was due to its involvement in a regulatory circuit with negative feedback loops and that this circuit could be removed without any apparent negative effects, if multiple genes were simultaneously deleted. Thus, this highlights how seemingly necessary but undesirable traits may be engineered as we unravel their systems-level context^46,47^, and that multiplex genome editing strategies^48^ and combinatorial genome editing screens^49,50^ will be particularly important in such research.

## ACKNOWLEDGEMENTS

This work was supported by the Novo Nordisk Foundation through the Technical University of Denmark (NNF16CC0021858, NNF10CC1016517, NNF20SA0066621) and from the NIGMS (R35 GM119850). NP and HFK further received funding from the European Union’s Horizon 2020 research and innovation programme under the Marie Skłodowska-Curie grant agreement No. 642663. EC is supported by the Österreichische Forschungsförderungsgesellschaft (FFG) grant number: 853223, Austrian Science Fund (FWF), grant numbers: P 32900 and P 33430 and Doc Fund DOC59-B33 and by a Christian Doppler Laboratory for recombinant protein production in mammalian cells.

## CONTRIBUTIONS

IMM carried out oxygen uptake experiments. IMM, NLC, and IMDM carried out the ^13^C experiments and data analysis. SMN generated protein producing pools. MD, KLJ, EMJ, and HH ran fed-batch experiments. JA, ZS, KKB, PM, SPB, NP, DL, HH, KJLCK, TK, JMCZ, SS, and SK generated and characterized cell lines. AGT and JM carried out phosphoproteomic analysis of Pdh. SK and HMP carried out protein purification. AHH carried out glycan analysis. AAM carried out the Seahorse experiment and analysis. EK and LS generated and characterized the Rituximab producing clones using BACs. AB and EC oversaw the BAC based cell line generation process. NB oversaw the Seahorse experiment. LKN oversaw the oxygen uptake and ^13^C experiments. HFK oversaw the generation of the Rituximab-producing clone (C6_2) and derivatives. GML oversaw pool generation. BGV oversaw cell line engineering. HH and NEL conceived and designed the study and wrote the manuscript.

## COMPETING INTERESTS

A patent based on this work has been issued with HH and NEL as inventors (US Patent 11,242,510, WO2017192437). AB and EC hold a patent about the BAC-based expression system (WO2010060844A1).

## METHODS

### Transfection

CHO-S^TM^ cells (Gibco Cat. # A11557-01) were transfected using either FreeStyle MAX reagent (Gibco Cat. # 16447100) or FuGENE HD reagent (Promega Cat. # E2311). For both procedures, at the day prior to transfection, viable cell density was adjusted to 8×10^5^ cells/mL in transfection medium: CD CHO medium (Gibco Cat. #10743-029) supplemented with 8 mM L-glutamine (Lonza Cat. # BE17-605F). With the FreeStyle MAX procedure, at the day of transfection, viable cell density was adjusted to 1×10^6^ cells/mL in an MD6 plate (Falcon Cat. # 351146) containing 3 mL transfection medium per well. For each transfection, 1.9 µg Cas9-2A-GFP plasmid DNA and 1.9 µg gRNA plasmid DNA was diluted in 60 µL OptiPro SFM (Gibco Cat. # 12309019). Separately, 3.8 µL FreeStyle MAX reagent was diluted in 60 µL OptiPro SFM and the two mixtures were incubated for 5 minutes at room temperature. After incubation, the plasmid DNA/OptiPro SFM mixture was added to the FreeStyle MAX/OptiPro SFM mixture and incubated at room temperature for an additional 20 minutes. The resultant 120 µL DNA/lipid mixture was added dropwise to the cells in one well. With the FuGENE HD procedure, at the day of transfection, viable cell density was adjusted to 8×10^5^ cells/mL in an MD6 plate containing 3 mL transfection medium per well. For each transfection, 1.5 µg Cas9-2A-GFP plasmid DNA and 1.5 µg gRNA plasmid DNA was diluted in 75 µL OptiPro SFM. Separately, 9.0 µL FuGene HD reagent was diluted in 66 uL OptiPro SFM. The plasmid DNA/OptiPro SFM mixture was added to the FuGENE HD/OptiPro SFM mixture and incubated at room temperature for 5 minutes and the resultant 150 µL DNA/lipid mixture was added dropwise to the cells in one well. Plasmids were constructed using the uracil-specific excision reagent (USER) cloning method as described previously^51^, with the sgRNA1_C plasmid as a backbone.

### Single cell sorting and expansion

Transfected cells were single cell sorted 48 hours post transfection, using the FACSJazz, based on green fluorescence with gating determined by comparison to non-transfected cells. Sorting was done into MD384 plates (Corning Cat. # 3542) containing CD CHO medium (Gibco Cat. # 10743-029) supplemented with 8 mM L-glutamine (Lonza Cat. # BE17-605F), 1% antibiotic-antimycotic (Gibco Cat. # 15240-062), and 1.5% HEPES buffer (Gibco Cat. # 15630-056). After 15 days, colonies were transferred to an MD96F plate (Falcon Cat. # 351172) containing CD CHO medium supplemented with 8 mM L-glutamine, 2 mL/L anti-clumping agent (Gibco Cat. # 0010057AE), and 1% antibiotic-antimycotic.

### Clone genotyping

After two days, 50 uL cell suspension from each well was transferred to a MicroAmp Fast 96 well reaction plate (Thermo Cat. # 4346907), along with 5×10^5^ wildtype cells as a control. The plate was centrifuged at 1000 x g for 10 minutes, then the supernatant was removed via rapid inversion. 20 µL of QuickExtract DNA Extraction Solution (Epicentre Cat. # QE09050) (prewarmed to 65°C) was added to each well and mixed via pipetting. The plate was then processed in the thermocycler (65°C for 15 minutes followed by 95°C for 5 minutes).

Amplicons were generated for each gene of interest per well using Phusion Hot Start II DNA Polymerase (Thermo Cat. # F549L) and verified to be present visually on a 2% agarose gel. Amplicons from each well had unique barcodes, allowing them to be pooled and purified using AMPure XP beads (Beckman Coulter Cat. # A63881) according to manufacturer’s protocol, except using 80% ethanol for washing steps and 40 uL beads for 50 uL sample. Samples were indexed using the Nextera XT Index kit attached using 2x KAPA HiFi Hot Start Ready mix (Fisher Scientific Cat. # KK2602). AMPure XP beads were used to purify the resulting PCR products.

DNA concentrations were determined with the Qubit 2.0 Fluorometer and used to pool all indices to an equimolar value and diluted to a final concentration of 10 nM using 10mM Tris pH 8.5, 0.1% Tween 20. The average size of the final library was verified with the Bioanalyzer 2100. The amplicon library was then sequenced on an Illumina MiSeq.

Insertions and deletions were identified by comparison of expected vs. actual amplicon size. Clones with frameshift indels in all alleles of the Ldha gene were selected for expansion.

### Batch culture

CHO-S^TM^ cells (Gibco Cat. # A11557-01) and derivative clones were cultured in CD CHO medium supplemented with 8 mM L-glutamine and 2 mL/L of anti-clumping agent. All cultures were maintained in an incubator at 37°C, 5% CO_2_, 70-95% humidity and 25 mm throw, while shaking at 120 rpm. Cell growth and viability were monitored using the NucleoCounter NC-200 Cell Counter (ChemoMetec, Denmark) based on two fluorescent dyes, Acridine Orange and DAPI for the total and dead cell populations, respectively. Metabolite concentrations were measured using the BioProfile 400 (Nova Biomedical, USA).

### Ldh enzymatic assay

Lactate dehydrogenase activity was measured using the Cytotoxicity Detection Kit PLUS (LDH) from Roche (Basel, Switzerland) according to the instructions of the manufacturer. In short, cell pellets (1×10^6^) were resuspended in 250 µl 1% Triton X-100, 50 mM TRIS-HCl pH 7.5 and lysed by incubation for 30 minutes at 37°C. All samples were clarified by centrifugation for 5 minutes (14290xg) at room temperature. Supernatants were diluted 27-fold with PBS and transferred to a 96 well plate for measurement.

### Western blotting

For Ldha (and LDHB in HEK) blots, cell pellets were resuspended in 50 mM Tris pH 8.0, 150 mM NaCl, 1 mM EDTA, 1% Triton X-100, 1 pill cOmplete/50mL (Roche) and lysed by incubation for 30 minutes on ice. Lysates were clarified via centrifugation at 14290 x g for 10 minutes at room temperature. After separation by SDS-PAGE, proteins were transferred onto PVDF membranes and probed with antibodies against Vinculin (V9131, 1:1000, Sigma, St. Louis, MO, USA) and LDHA (#2012, 1:1000, Cell Signaling Technology, Danvers, MA, USA) or LDHB (ab53292, 1:50000, Abcam). Proteins of interest were detected with Cy5 and Cy3 conjugated secondary antibodies (29-0382-78 and 29-0382-75, 1:2500, Amersham, Little Chalfont, UK) and visualized on an AI600 imager (Amersham). Quantification was done using ImageQuant TL 1D v8.1. For nonfluorescent blots, samples were treated as above except a nitrocellulose membrane was used, the antibody against Vinculin was diluted 1:2000, HRP-conjugated donkey anti-rabbit (ab6802, 1:5000, Abcam, Cambridge, UK) or goat anti-mouse (P0447, 1:5000, Dako) were used as secondary antibodies, and detection was with the Amersham ECL detection system. All dilutions, washing, and blocking steps were done with PBST + 0.1% Tween-20 + 5% skim milk.

For Pdh blots, cell pellets were resuspended in 50 mM Tris pH 8.0, 150 mM NaCl, 1 mM EDTA, 1% Triton X-100, 1x Halt Protease & Phosphatase Inhibitor (Thermo Fisher Scientific Cat. # 78442). After separation by SDS-PAGE, proteins were transferred onto nitrocellulose membranes and probed with antibodies against Pdh (ab168379, 1:1000, Abcam, Cambridge, UK), phospho-(S293)-Pdh (ab92696, 1:1000, Abcam, Cambridge, UK), and Vinculin (as above). HRP-conjugated donkey anti-rabbit (ab6802, 1:5000, Abcam, Cambridge, UK) or goat anti-mouse (P0447, 1:5000, Dako) were used as secondary antibodies, and detection was with the Amersham ECL detection system. Dilution, washing, and blocking was done with TBST + 0.1% Tween-20 + 5% BSA.

### MS phosphoproteomic analysis of Pdh

Resuspended cell pellets taken for Pdh were further processed to remove Triton X-100 and enrich Pdh. Triton X-100 removal was carried out on 100 uL suspension with Pierce^TM^ Detergent Removal Spin Columns (Thermo Fisher Scientific Cat. # 87777) following the manufacturer’s protocol and using 50 mM Tris pH 8.0, 150 mM NaCl, 1 mM EDTA as the wash/equilibration buffer. The eluate was resuspended in a total volume of 1 mL 50 mM Tris pH 8.0, 150 mM NaCl, 1 mM EDTA with 3 uL of anti-Pdh (ab168379, Abcam) and was incubated overnight at 4°C with rotation. The following day 50 uL 50% protein A sepharose (previously washed 3x with 50 mM Tris pH 8.0, 150 mM NaCl, 1 mM EDTA) was added and samples incubated for 2 hours at 4°C with rotation. After incubation, samples were washed 3x with 1 mL 50 mM Tris pH 8.0, 150 mM NaCl, 1 mM EDTA prior to elution with 50 uL 0.1M sodium citrate buffer pH 3.0 which was subsequently neutralized with 50 uL 50 mM Tris pH 8.0, 150 mM NaCl, 1 mM EDTA and frozen prior to further analysis.

#### Trypsin digestion and peptide desalting

Protein samples were resuspended in 50 mM ammonium bicarbonate and reduced with 5LmM Tris(2-carboxyethyl)phosphine hydrochloride, pH 7.0 for 45 min at 37°C, and alkylated with 25LmM iodoacetamide (Sigma) for 30 min at RT. Samples were digested with trypsin (1/100, w/w, Sequencing Grade Modified Trypsin, Porcine; Promega) overnight. Digestion was stopped using 10% trifluoracetic acid (Sigma). Peptide purification was performed by C-18 AssayMAP Bravo (Agilent), according to the manufacturer’s guideline. Samples were lyophilized and resuspended in 50Lμl HPLC-water (Fisher Chemical) with 2% acetonitrile and 0.2% formic acid (Sigma) before LC-MS/MS analysis.

#### LC-MS/MS for proteomics analysis

Peptide and Phosphopeptide analyses were performed on a Q Exactive HF-X mass spectrometer (Thermo Fisher Scientific) coupled to an EASY-nLC 1200 ultra-HPLC system (Thermo Fisher Scientific). Peptides were trapped on precolumn (PepMap100 C18 3 μm; 75LμmL×L2Lcm; Thermo Fisher Scientific) and separated on an EASY-Spray column (ES903, column temperature 45L°C; Thermo Fisher Scientific). Mobile phases of solvent A (0.1% formic acid), and solvent B (0.1% formic acid, 80% acetonitrile) were used to run a linear gradient from 5% to 38% over 90 min at a flow rate of 350Lnl/min. Data was acquired in the MS instrument in data-dependent mode according to the manufacturer’s default for ‘high sample amount’. Raw files were searched using Proteome Discoverer 2.5 and against the UniProt human database. The precursor and fragment mass tolerances were set to 10 ppm and 0.02LDa, respectively. Methionine oxidation and serine/threonine/tyrosine phosphorylation were set as dynamic modifications. Carbamidomethylation of cysteines was included as a static modification.

### Bioreactor cultures for determination of oxygen uptake

WT and 0-F5 cells (n=2 per line) were grown in CD OptiCHO medium (Gibco Cat. # 12681011) containing 4 mM glutamine, 50 ppm Antifoam C (Sigma A8011), 1.05 mL/L Anti-clumping agent, and 10.5 mL/L antibiotic-antimycotic. Cultures were grown in DASGIP parallel bioreactors (single-use, Eppendorf) with a starting volume of 270 mL at a temperature of 37°C, agitated at 200 rpm using one pitched blade impeller. The pH was maintained continuously at 7.10±0.02 with 1 M sodium bicarbonate or CO_2_. Dissolved oxygen was maintained at 40% using pure oxygen, air, or a mixture, using an aeration flow fixed at 3 L·h^-1^. Glutamine was added on demand to be maintained between 1-4 mM. Cultures were sampled every 6-8 hours to determine cell count and viability via NucleoCounter NC-200 Cell Counter (ChemoMetec) and targeted metabolites via BioProfile 400 (Nova Biomedical).

Volumetric oxygen uptake rate (vOUR) was determined via the dynamic method^52–54^. Briefly, every 6-8 hours dissolved oxygen (DO) levels were increased to 60%. Afterwards, DO control and gas supply via sparger were paused while an N_2_ flow (0.07 vvm) was introduced into the bioreactor headspace to obtain an oxygen-free gas phase and avoid oxygen transport back to the culture medium. OUR calculations were performed by monitoring the DO profile between 60 and 30% to quantify the respiratory activity of the cells while accounting for oxygen desorption from the liquid to gas phase using an experimentally determined desorption constant, K_des_ (Equation 3). DO control resumed when levels dropped below 30%. K_des_ is highly dependent on the culture volume so different measurements were performed in triplicate prior to inoculation in the working volume range (270-170 mL). The real K_des_ was then used depending on the real culture volume to determine vOUR through equation 3.

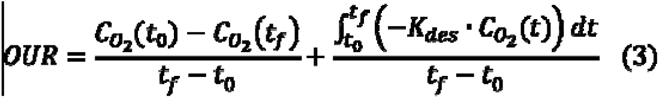

Specific oxygen consumption rate (q_Oxygen_) was calculated by fitting a regression line with zero intercept (vOUR and VCD proportional) allowing for a variation in slope due to the cell type to data points between t=20 and 68 hrs. More explicitly, in R, lm(formula = OUR ∼ -1 + VCD + VCD:celltype). The calculated slope represents the cell specific oxygen uptake rate.

### Transcriptomic, endometabolomic, and ^13^C experiments

WT and 0-F5 cells (n=4 per line) were grown in bioreactors as above. When cell density reached ∼3×10^6^ cells/mL, samples were taken for transcriptomics and endometabolomics from 3 bioreactors per line. For transcriptomics, approximately 5×10^6^ cells were removed from the bioreactor and centrifuged at 200 x g for 2 min (VWR). The supernatant was discarded and the cell pellet was frozen in liquid nitrogen and stored at -80°C for further analysis.

The protocol for quenching and extraction for endometabolomics was based on existing methods^55^. Approximately 25×10^6^ cells were drawn from the bioreactor into a 50mL tube containing 40 mL of ice cold 0.9% (w/v) NaCl. The tube was centrifuged at 1000 x g for 1 min at 0.5°C. Supernatant was discarded and the cell pellet was resuspended in 50 mL of ice cold 0.9% NaCl for washing. The tube was again centrifuged at 1000 x g for 1 min at 0.5°C. Supernatant was discarded and the cell pellet was resuspended in 50% aqueous acetonitrile solution and incubated for 10 min at 0°C including several rounds of vortexing (a volume of 1 mL extraction solution is used per 5×10^6^ cells). The tube was centrifuged at 7100 rcf for 25 min at 5°C. The supernatant was transferred to a new 15 mL tube, frozen on dry ice, freeze dried and stored at -80°C. Prior to analysis, the lyophilized metabolites were resuspended in MilliQ purified water. The intracellular metabolite concentrations were analyzed using LC-MS following a previously described protocol^56^.

At the same time point, a mix of 90% U-Glc and 10% 1-^13^C-Glc ([Glc]=500 mM) (Eurisotop) was added to the remaining bioreactor for each line (to increase the glucose concentration by ∼20 mM). Glutamine was also added to increase the concentration up to 8 mM to avoid adding more glutamine during the time experiment. From this point, samples were taken every 2 hours for 20 hours for endometabolomics.

Endometabolomic samples for the ^13^C phase were analyzed with GC-MS. GC-MS analysis was performed using a Bruker 436 Gas Chromatography System equipped with a BPX5 column, 30m x 0.25mm, 0.25 µm (SGE Analytical Science) coupled to a Scion QqQ mass spectrometer (Bruker) operating in El mode (70eV). Samples were derivatized with N-Methyl-N-trimethylsilyl trifluoroacetamide and 1 µL was injected into a 280°C injection port in split mode (1:4). Helium flow was maintained at 1 mL/min. The column temperature was held at 70°C for 4 min, ramped at 15°C/min to 310°C and held for 20 min. After a 5 min solvent delay, the mass spectrometer collected data in full scan mode between 45-750 Da.

For each molecule, different fragments were generated in the derivatization protocol with different C information, as described previously^57^. The peak areas ratio of non-labelled and labeled molecules were calculated for the different fragments using Xcalibur^TM^ Software (Thermo Scientific). A correction for naturally occurring isotopes was performed using MIDcor^58^. The labeling incorporation for each metabolite was determined by calculating the ratio of the sum of all the labeled isotope ratios and the non-labeled molecule at each time point.

Separately, WT and 0-F5 cells (n=4 per line) were cultured in the ambr^®^15 (Sartorius, Germany) system in CD CHO medium supplemented with 0.01% antibiotic-antimycotic and 8 mM glutamine. The cultures were inoculated at 0.5×10^6^ cells/mL with an initial volume of 13.5 mL.

Dissolved oxygen was maintained using a mixture of air and oxygen gas with a set point of 40%. The temperature was set at 37 °C and the stirring at 900 rpm. 20 μL of 5% Antifoam C Emulsion was added daily to avoid foam accumulation.

To generate the time course for the ^13^C analysis approximately 16 mM of U-^13^C-Glc (Eurisotop) was added to the bioreactors after cell density reached ∼3×10^6^ cell/mL. Endometabolomics samples were then taken at 0, 2, 4, 8 and 14 hours after spiking with the labeled glucose. The endometabolomics samples were collected by aliquoting 1 mL (∼3×10^6^ cells) of the culture into a 5 mL-tube containing 4 mL of ice cold 0.9% (w/v) NaCl. After, the sample was centrifuged at 1000 rcf at -5 °C for 1 minute. The supernatant was then discarded and 2.5 mL more of ice cold NaCl solution that was mixed by inversion to resuspend the pellet. Following, the sample was centrifuged again at 1000 rcf at -5 °C for 1 minute and again the supernatant was discarded. The samples were then snap frozen and stored at -80 °C prior to analysis.

The cell pellets were extracted as previously described^55^. 700 µL of an extraction buffer containing Acetonitrile:Methanol:Water in a ratio of 2:2:1 (v/v/v) as well as 0.1% Formic acid (v/v). After resuspending the thawed pellets, the samples were transferred to 2-mL tubes and kept on ice. Add 100 µL of extraction buffer to the tubes and vortexed to mix. The samples were then incubated at -80 °C for 20 minutes. The centrifuge was pre-cooled for 20 minutes at -9 °C before centrifuging the samples at 17000 × g for 5 minutes. The supernatant was transferred to a new tube and stored at -80 °C. Afterwards, the pellets were washed in 200 µL extraction buffer, vortexed and incubated at -80 °C for 20 minutes before centrifuging at 17000 × g for 5 more minutes with the supernatant being transferred to the same tube as before. The washing process was repeated one more time, resulting in a total sample volume of 1200 µL for each tube. The samples were dried at 30 °C for approximately 8 hours before being stored at -80 °C until analysis.

Metabolomics samples from the ambr^®^15 were then analyzed using a reverse-phase ion-pairing-liquid chromatography-tandem mass spectrometry (LC-MS/MS) method performed using a Sciex QTrap 5500 run in MRM mode, as well as a Shimadzu LC-20/30 series LC system with a Waters XSelect HSS T3_2.5µm _2.1×150mm_XP column. Acquisition was performed as previously described^56^. Each injection was 10 µL and quantification was performed using SciexOS software with relative isotope abundances estimated based on peak area.

The enrichment of TCA metabolites was measured by normalizing the sum of the labeled isotopes to that of the extracellular glucose labeling fraction, as determined by the relative amount of U-^13^C-Glc as measured by GC-MS.

### Transcriptomic sequencing and analysis

The protocol for RNA extraction and quantification (about 20 million reads per sample) has been described previously^59^. Reads were trimmed using trimmomatic^60^, aligned against the Chinese hamster reference genome (RefSeq assembly accession: GCF_003668045.3) and Chinese hamster mitochondrial reference genome (RefSeq: NC_007936.1) using STAR^61^, and quantified with htseq^62^. Differential expression analysis was done using DESeq2^63^ and genes with an adjusted p-value<0.05 were passed to Ingenuity Pathway Analysis^64^ (QIAGEN Inc.) and used in the Expression Analysis module to identify perturbed canonical pathways based on directional change of gene expression.

### Seahorse assay

0-F5 and *Lama2* knockout CHO-S cells (which are not expected to have an altered metabolic phenotype) were thawed and maintained in CD CHO medium supplemented with 8 mM glutamine (Thermo-Fisher Scientific, Cat. # 25030024) and 1:500 PVA (Acros Organics, Cat. # 180300010) as an anti-clumping agent. After recovery, cells were counted using a Guava benchtop flow cytometry and stained using Guava ViaCount, (Luminex, Cat. # 4000-0040), centrifuged at 1000 x g for 5 minutes and resuspended in 2 mL of assay media (Seahorse XF base medium without phenol red (Agilent, Cat. # 103334-100), 1 mM pyruvate (Acros Organics, Cat. # 132151000), 2 mM glutamine, 10 mM glucose (Agilent, Cat. # 103577-100), 5 mM HEPES (Agilent, Cat. # 103337-100 ). Cells were plated in Seahorse 96 well plates (Agilent, Cat. # 103729-100) previously coated with Corning Cell Tak cell adhesive (Corning, Cat. # 10317081) following manufacturer’s instructions and analyzed using the Seahorse XF Glycolytic Rate Assay (Agilent, Cat. # 103344-100) following manufacturer’s instructions. Each cell type was grown in 2 independent flasks and each flask was seeded into 23 wells at an initial density of 20000 cells/well.

### BAM15 toxicity assay

A stock solution was made by mixing 5 mg of BAM15 (Sigma Aldrich, Cat. # SML1760-5MG) with 2.17 mL of DMSO (∼6.77 mM). This was diluted in fresh CD CHO medium containing 8 mM glutamine and 2 mL/L anti-clumping agent to a concentration of ∼2.25 µM (4.33 uL stock solution into 13 mL medium).1.5 mL of this solution was mixed with 1 mL cell suspension taken from cells in mid exponential growth (∼1×10^6^ cells/mL) and aliquoted into 6 well plates for analysis (final concentration ∼1.35 µM BAM15). All cells were grown in duplicate and measured daily for cell count and viability using the NucleoCounter NC-200.

### Shake flask culture characterization of full Pdk + Ldha knockout CHO-S clones for transcriptomic characterization

Clones 0-B6, 0-F4, 0-F5 and the wildtype CHO-S cell line were grown in batch culture. On day 3, 2×10^6^ cells were harvested for RNA-Seq. Cells were centrifuged at 200 x g for 2 min, the supernatant was discarded and the cell pellet was frozen in liquid nitrogen and stored at -80°C until processed.

### Cell line generation-Rituximab producing pools

Host cell lines were transfected with a vector containing the GS and antibody genes using FreeStyle MAX (Invitrogen) according to the manufacturer’s protocol. Twenty-four hours after transfection, cells were inoculated at a concentration of 1×10^6^ cells/mL into 125-mL Erlenmeyer flasks containing 50 mL of selection medium (CD CHO supplemented with GSEM, 25 µM MSX, 300 µg/mL zeocin and 2 µL/mL anti-clumping agent). The flasks were subsequently incubated in a Climo-shaking incubator (Kuhner AG, Basel, Switzerland) at 110 rpm in a humidified 5% CO_2_/air mixture at 37°C. Every third day, medium was exchanged following centrifugation of cells at 1200 rpm for 5 minutes and reseeded at up to 1×10^6^ cells/mL. When viable cell concentration first exceeded 2×10^6^ cells/mL, cells were reseeded at 5×10^5^ cells/mL; each subsequent reseeding was at 3×10^5^ cells/mL. When viability exceeded 90%, pool selection was considered complete.

### Cell line generation-Other proteins

As above except 24 hours after transfection, inoculation was into 10 mL CD CHO supplemented with GSEM, 25 µM MSX, and 2 µL/mL anti-clumping agent. After 9 and 15 days of selection, cells were subjected to increased selective pressure (100 and 200 µM MSX, respectively). After 34 days of selection, cells were single cell sorted using the FACSJazz in MD384 plates containing CD CHO medium supplemented with 8 mM L-glutamine, 1% antibiotic-antimycotic, and 1.5% HEPES buffer. Enbrel-transfected pools were first surface stained to enrich for high producers^65^. After 14 days, clones were transferred to MD96F plates containing CD CHO medium supplemented with 8 mM L-glutamine, 1% antibiotic-antimycotic, and 2 mL/L anti-clumping agent. After 3 days of growth in MD96F plates, confluence was assessed using the Celígo Imaging Cell Cytometer (Nexcelom Bioscience, Lawrence, MA) with the label-free, bright field confluence application and titer quantified via ELISA (GDF5) or Octet (all other proteins).

### Cell line generation and protein quantification-The Antibody Lab

Cell lines expressing Rituximab and Herceptin were generated using the BESTCell^TM^ technology as described previously^66^. In brief, expression constructs were generated using BAC (bacterial artificial chromosomes) vectors and transfected using Amaxa Nucleofector Kit V (Lonza; VCA-1003). Upon selection using neomycin (1 to 1.5 mg/mL), surviving single cells were isolated using FACS sorting. After colonies had formed, the number of BAC integrations was determined using qPCR and secretion of the recombinant protein into the supernatant was confirmed using an in-house ELISA assay. The top clones in terms of transgene copy number were further evaluated by fed-batch in 24 deep-well plates using ActiPro medium and Cell Boost 7a and 7b as feeds. Final assessment of quantity and quality of the recombinant protein in supernatant was done using non-reducing gel electrophoresis (Thermo Fisher Scientific) combined with coomassie staining and near-infrared fluorescent imaging (Li-Cor).

### Protein quantification

#### Rituximab, Enbrel, Erythropoietin, and C1 esterase inhibitor

These four proteins were quantified using biolayer interferometry with an Octet RED96 (Pall Corporation, Menlo Park, CA). Octet System Data Analysis 7.1 software was used to calculate binding rates and absolute concentrations. Specific assay setups are described below.

##### Rituximab/Enbrel

ProA biosensors (Fortebio 18-5013) were hydrated in PBS and preconditioned in 10 mM glycine pH 1.7. A calibration curve was prepared using human IgG at 400, 200, 100, 50, 25, 12.5, and 6.25 mg/L serially diluted in CD CHO medium. Sample supernatants were collected after centrifugation and association was performed for 120 seconds with a shaking speed of 200 rpm at 30°C. A separate calibration curve was used for the ambr15 experiments (Figure 5 and Supplementary Figure 15) using purified Rituximab at 400, 200, 100, 50, 40, 25, 20, 12.5, 10, 6.25, 5, 2.5, 1.25, and 0.625 mg/L serially diluted (from 400 and 40 mg/L stocks) in ActiPro medium to account for matrix effects.

##### Erythropoietin/C1 esterase inhibitor

Established quantification protocols were followed^67,68^. Briefly, streptavidin biosensors (Fortebio 18-5021)—following hydration in PBS—were functionalized with CaptureSelect biotin anti-EPO conjugate or CaptureSelect biotin anti-C1INH conjugate (Thermo Fisher Scientific) at 5 µg/mL in PBS, and blocked in PBS containing 1 µg/mL biocytin (600 and 300 s incubation steps, respectively), preceded by and followed by hydration in PBS for 60 s. Standards (EPO: GenScript Z02975-50, C1INH: R&D Systems 2488-PI-200) were prepared in spent medium. Standards and samples were diluted two-fold. The dilution buffer consisted of water containing 0.1% BSA, 0.1% Tween20, 500 mM NaCl, and—for EPO only—20 mM, pH 4 citric acid. After equilibration in spent medium for 300 or 60 s (EPO or C1INH, respectively), samples and standards were measured for 200 s with a shaking speed of 1000 rpm at 30 °C. Tips were regenerated after each measurement in 10 mM, pH 12 Na_2_PO_4_ (EPO) or 2 M MgCl2 50 mM, pH 7.5 TRIS HCl and neutralized in PBS.

##### GDF5

Titers were quantified using ELISA (R&D Systems Cat. # DY853-05) following manufacturer’s protocol.

### Fed-batch culture

#### Clone P-F5

Cells were inoculated at a density of 3×10^5^ cells/mL in ambr15 bioreactors with a starting volume of 15 mL at a temperature of 37°C, agitated at 900 rpm using pitched blade impellers. The pH was maintained at 7.1±0.05 with 1 M sodium bicarbonate or CO_2_ and calibrated every third day. Cells were sampled daily for viable cell count and major metabolites on the BioProfile FLEX2 and, starting on day 7, for Rituximab quantification using the OctetRED96. A variety of process parameters were tested (see Supplementary Note for all parameters varied). Below are the details for the vessel shown in Figure 5A-C.

Dissolved oxygen was maintained at 50% using pure oxygen, air, or a mixture, as needed. The agitation rate was adjusted as needed to maintain proper oxygenation. The temperature was shifted to 33° on day 6.

Cells were grown in ActiPro medium (GE Healthcare Cat. # SH31039.02) supplemented with 4% GlutaMAX (Thermo Fisher Cat. # 35050061), 1% antibiotic-antimycotic, and 2 mL/L anti-clumping agent. Antifoam C was added as needed over the course of culture. Cell Boost 7a and 7b (GE Healthcare) were added from day 3-7 (inclusive) as a percentage of culture vessel volume:

Day 3: 3% 7a, 0.3% 7b
Day 5: 4% 7a, 0.4% 7b
Day 6: 4% 7a, 0.4% 7b
Day 7: 4% 7a, 0.4% 7b

Cell Boost 7a was added as 3 separate boluses.

Starting day 9, cells were fed 3% vessel volume 1x Efficient Feed C+ (Thermo Fisher Cat. # A2503101) (split into 3 separate boluses) instead of Cell Boost 7a/7b. Glucose was fed daily (split into 2 separate boluses) to a setpoint of 28 mM as needed using a 2222 mM glucose solution. Cells received 1.5% vessel volume GlutaMAX on day 5 and 8 and 1% vessel volume 1 mM thiamine on day 8.

### Introducing the Warburg-null phenotype into a Rituximab-producing clone

Plasmid used for generation of C6_2 cell line consisted of the Rituximab expression cassette (CMV-HC-BGHpA-EF1α-LC-BGHpA) and the vector backbone with the selection marker (SV40-NeoR-SV40pA). CHO-S cells (Thermo Fisher Scientific) were transfected with Rituximab plasmid using FuGENE®□ HD transfection reagent (Promega), followed by G418 selection (500 µg/mL; Sigma Aldrich). After stable polyclonal pools were established, single-cell cloning, clone screening for high-producers, titer and specific productivity measurement were performed as described in our previous study^65^. C6_2 was selected based on high titer and was tested for stability over two months.

C6_2 was transfected with gRNAs targeting the Pdks and Ldha. As a control for the transfection process, a second population was transfected with only the plasmid containing Cas9. Single-cell sorting and expansion was carried out as for non-producing CHO-S cells. Warburg-null clones were selected based on no lactate production as determined by the BioProfile 400. Five control clones were selected randomly for expansion.

#### Clone P-H6

Cells were inoculated at a density of 2.5×10^5^ cells/mL in DASGIP bioreactors with a starting volume of 270 mL at a temperature of 37°C, agitated at 200 rpm using pitched blade impellers. The pH was maintained continuously at 7.10±0.02 with 1 M sodium bicarbonate or CO_2_. Dissolved oxygen was maintained at 40% using pure oxygen, air, or a mixture, as needed.

Cells were grown in ActiPro medium supplemented with 8 mM L-glutamine, 1% antibiotic-antimycotic, and 1 mL/L anti-clumping agent. Antifoam C was added as needed over the course of culture. Glucose was maintained between 10-24 mM. Glutamine was maintained between 2-4 mM. Cell Boost 7a and 7b (GE Healthcare) were added starting at day 4 at roughly the following volumes:

Day 4: 4 mL 7a + 0.4 mL 7b
Day 5: 8 mL 7a + 0.8 mL 7b
Day 6: 12 mL 7a + 1.2 mL 7b
Day 7: 17.5 mL 7a + 1.6 mL 7b

A 2222 mM glucose solution and a 200 mM glutamine solution were used to control glucose and glutamine levels at the desired levels via a once-daily feeding based on the measured concentration, doubling time, and calculated consumption rates. Cultures were sampled daily for cell growth and viability using the NucleoCounter NC-200 Cell Counter, metabolite concentrations using the BioProfile 400, and rituximab concentration using the OctetRED96.

### Glycan analysis

After centrifugation and filtration to remove the cells and cell debris, the secreted mAbs were purified from 5 mL aliquots of culture supernatant. Purification was done using protein A affinity chromatography (MabSelect recombinant protein A agarose, GE Healthcare, Uppsala, Sweden), according to the manufacturer’s protocol. Purified mAbs were fluorescently labeled with GlykoPrep Rapid N-Glycan kit (ProZyme, Hayward, CA), according to the manufacturer’s protocol. N-linked glycan analysis was performed by LC-MS system using a Thermo Ultimate 3000 HPLC with the fluorescence detector coupled on-line to a Thermo Velos Pro Iontrap MS, as described previously^48^.

### Warburg-null HEK 293

A similar protocol (to the one used for CHO lines) was used for HEK 293 cell line generation. Briefly, HEK 293F cells (Thermo Cat. # R79007) were maintained in FreeStyle 293 medium (Thermo Cat. # 12338018) supplemented with 2 mL/L anti-clumping agent and 1% antibiotic-antimycotic. Transfection with all 6 gRNAs (PDK1-4, LDHA, and LDHB, Supplementary Table 7) was carried out following the FreeStyle MAX transfection protocol used for CHO-S cells, except in FreeStyle 293 medium containing 1% antibiotic-antimycotic. Single cell sorting was into MD384 plates containing FreeStyle 293 medium + 1% antibiotic-antimycotic + 1.5% HEPES buffer + 1x InstiGRO HEK (Solentim Cat. # RS-1305). After 16 days, clones were moved to MD96F plates, containing FreeStyle 293 medium supplemented with 2 mL/L anti-clumping agent and 1% antibiotic-antimycotic. The same medium was used for subsequent expansion. A second round of transfection, single cell sorting, and expansion was carried out using the same protocol on intermediate clones (HEK-IM A and HEK-IM B) for genes that were not knocked out in the first round.

Genotyping was carried out as described for CHO-S for the first round of transfections (resulting in HEK-IM A and HEK-IM B). Final Warburg-null clones (HEK-WN) amplicons were prepared the same way but sequenced by Novogene with indel analysis via CRISPResso2^69^.

For batch culture, HEK 293F and derivative clones were cultured in FreeStyle 293 expression medium supplemented with 2 mL/L anti-clumping agent and 1% antibiotic-antimycotic. All cultures were maintained in an incubator at 37°C, 5% CO2, 70-95% humidity and 25 mm throw, while shaking at 120 rpm. Cell growth and viability were monitored using the NucleoCounter NC-250 Cell Counter (ChemoMetec, Denmark) based on two fluorescent dyes, Acridine Orange and DAPI for the total and dead cell populations, respectively. Metabolite concentrations were measured using the BioProfile FLEX2 (Nova Biomedical, USA).

### Warburg-null CHO-K1

Cells were transfected following the same protocol as CHO-S but single-cell sorted into 96 well U-bottom plates containing Ex-Cell CHO medium (Sigma Aldrich, Cat. # 14361C) supplemented with 4 mM glutamine, 2 g/L rHSA (Sigma, Cat. # A7223-1g), 5 mg/L rHTR (STEMCELL Technologies, Cat. #9655), 0.1% anti-oxidant (Sigma, Cat. # A1345-5mL), 1% antibiotic-antimycotic, and 1.5% HEPES. While cells were expanding, the medium was replaced slowly with standard growth medium (CD CHO supplemented with 8 mM glutamine, 1% antibiotic-antimycotic, 2 mL/L anti-clumping agent). Genotyping was carried out as described for CHO-S.

Batch culture was conducted in triplicate identically to CHO-S cultures. Cells were sampled daily for growth and viability using the NucleoCounter NC-200 Cell Counter and for metabolites starting on day 1 using the BioProfile FLEX2. On day 2 and 3, 1×10^6^ cells were harvested for RNA-Seq. Samples were centrifuged at 200 x g for 3 minutes, supernatant removed, and pellets frozen in liquid nitrogen and stored at -80°C until processed.

### Data Availability

RNA-Seq data is available on SRA (PRJNA746067). Mass spectrometry data files have been deposited in the MassIVE repository with the following identifier: MSV000095049. The raw data supporting the findings of this study are available from the corresponding author upon request.

