## Supplementary Information for "Multiplex genome editing eliminates the Warburg Effect without impacting growth rate in mammalian cells"

**Supplementary Figures and Tables**

*Supp. Figure 1: Maximum lactate concentrations for cultures in this study*

*Supp. Figure 2: Ldh blots and lactate levels*

*Supp. Figure 3: Warburg-null CHO-K1 batch culture*

*Supp. Figure 4: Warburg-null HEK 293F batch culture*

*Supp. Figure 5: pPdh and Pdh quantified*

*Supp. Figure 6: pPdh and Pdh blots*

*Supp. Figure 7: ^13^C labeling reveals increased TCA cycle activity in Warburg-null cells*

*Supp. Figure 8: Growth profiles of Warburg-null CHO-S cell lines in shake flask batch culture*

*Supp. Figure 9: Elimination of the Warburg effect results in cells with a prolonged growth phase that do not require base addition to maintain culture pH*

*Supp. Figure 10: Time-course profiles of essential amino acids in fed-batch culture*

*Supp. Figure 11: Eliminating the Warburg effect does not impact polyclonal pool generation*

*Supp. Figure 12: Outlier in pool generation process for clone 0-B6*

*Supp. Figure 13: Warburg-null clones can be used for production of diverse biotherapeutic proteins*

*Supp. Figure 14: Fed-batch screening product titers*

*Supp. Figure 15: Warburg-null cells producing Rituximab in ambr15 fed-batch*

*Supp. Figure 16: Batch-culture characterization and media optimization of clones derived from rituximab-producing line C6_2, following single-cell sorting*

*Supp. Figure 17: Rituximab glycoprofiles during fed-batch culture of parental and engineered cell lines*

*Supp. Table 1: Oligos used for cloning gRNAs and primers for MiSeq analysis of CHO genes targeted in this study*

*Supp. Table 2: Genotypes of isolated CHO Warburg-null clones*

*Supp. Table 3: Detailed genotypes of isolated CHO Warburg-null clones*

*Supp. Table 4: Source for gene expression data in Figure 1*

*Supp. Table 5: Genotypes of the isolated CHO-K1 clone*

*Supp. Table 6: Detailed genotype of the isolated CHO-K1 clone*

*Supp. Table 7: Oligos for cloning gRNAs and MiSeq primers for HEK lines*

*Supp. Table 8: Genotypes of isolated HEK clones*

*Supp. Table 9: Detailed genotypes of isolated HEK clones*

*Supp. Table 10: Mass spec based quantification of Pdh phosphorylation*

**Supplementary Note: Fed-batch culture**

**Supplementary Figures and Tables**

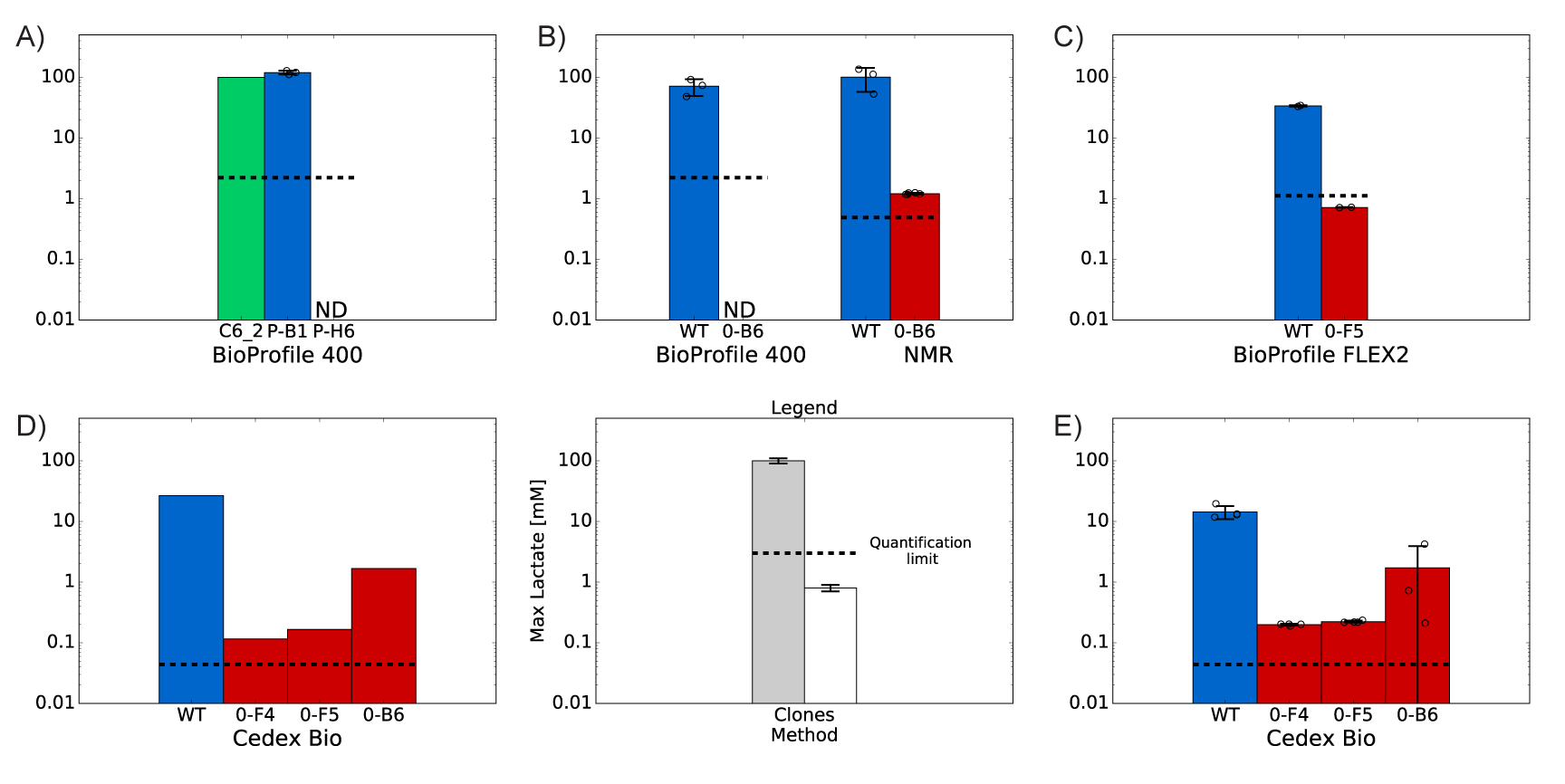

***Supplementary Figure 1: Maximum lactate concentrations for cultures in this study***

*Maximum lactate concentrations from fed-batch of (A) rituximab-producing cells (see also Figure 5D), (B) non-producing clone 0-B6 (see also Supplementary Figure 9), (C) non-producing clone 0-F5 (see also Supplementary Figure 9), and batch culture of (D) non-producing clones 0-B6, 0-F4, and 0-F5 (n=1 shake flask for each clone), or (E) rituximab producing pools derived from 0-B6, 0-F4, and 0-F5 (see also Supplementary Figure 11). Method of measuring lactate is shown below each plot with the quantification limit denoted by a horizontal dashed bar. ND: not detected (reported lactate 0 mM). Quantification limits for the methods are as follows: BioProfile 400-2.22 mM, BioProfile FLEX2-1.11 mM, NMR-0.49 mM, Cedex Bio-0.044 mM. Data shown as mean ± standard deviation.*

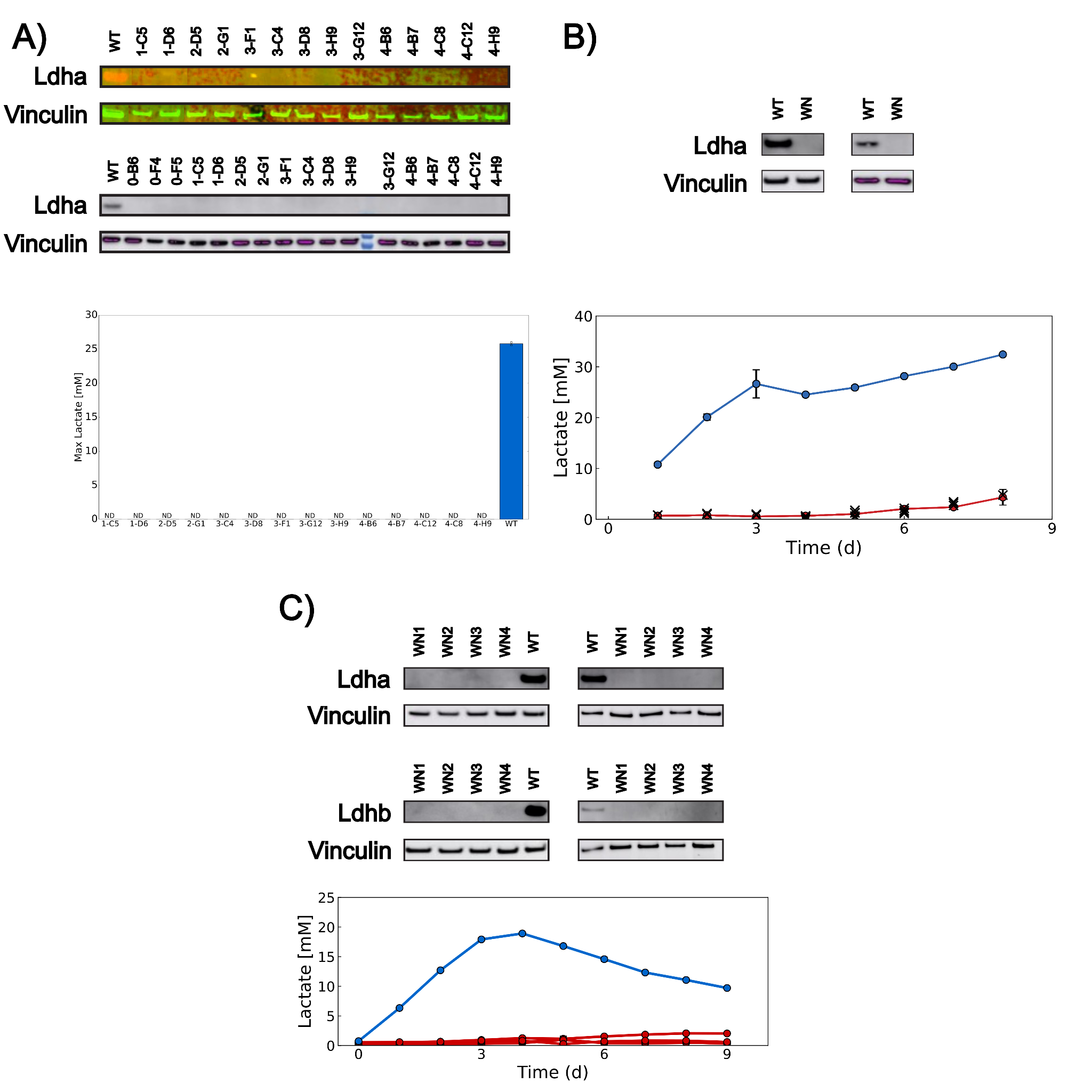

***Supplementary Figure 2: Elimination of the Warburg effect enabled by knockout of lactate dehydrogenase(s) in several mammalian cell lines***

*Targeting Pdks alongside Ldh(s) in (A) CHO-S, (B) CHO-K1, and (C) HEK 293F cells results in isolated clones with no detectable lactate dehydrogenase (duplicate blots shown) and negligible lactate production (n=2 shake flasks per cell line in (A), n=3 shake flasks per cell line in (B) and (C)). In all panels, wildtype cells are shown in blue, Warburg-null cells in red. Growth and glucose uptake rates for lactate levels in (A) shown in Figure 2A/B. Additional culture information for (B) and (C) shown in Supplementary Figure 5 and Supplementary Figure 6, respectively. Lactate levels (bottom plots) measured by BioProfile 400 in (A) and BioProfile FLEX2 in (B) and (C). In (B), black Xs mark measurements for blank solution. ND: not detected. Detailed cell line genotypes for CHO-S, CHO-K1, and HEK 293F clones are shown in Supplementary Tables 2/3, 5/6, and 8/9, respectively.*

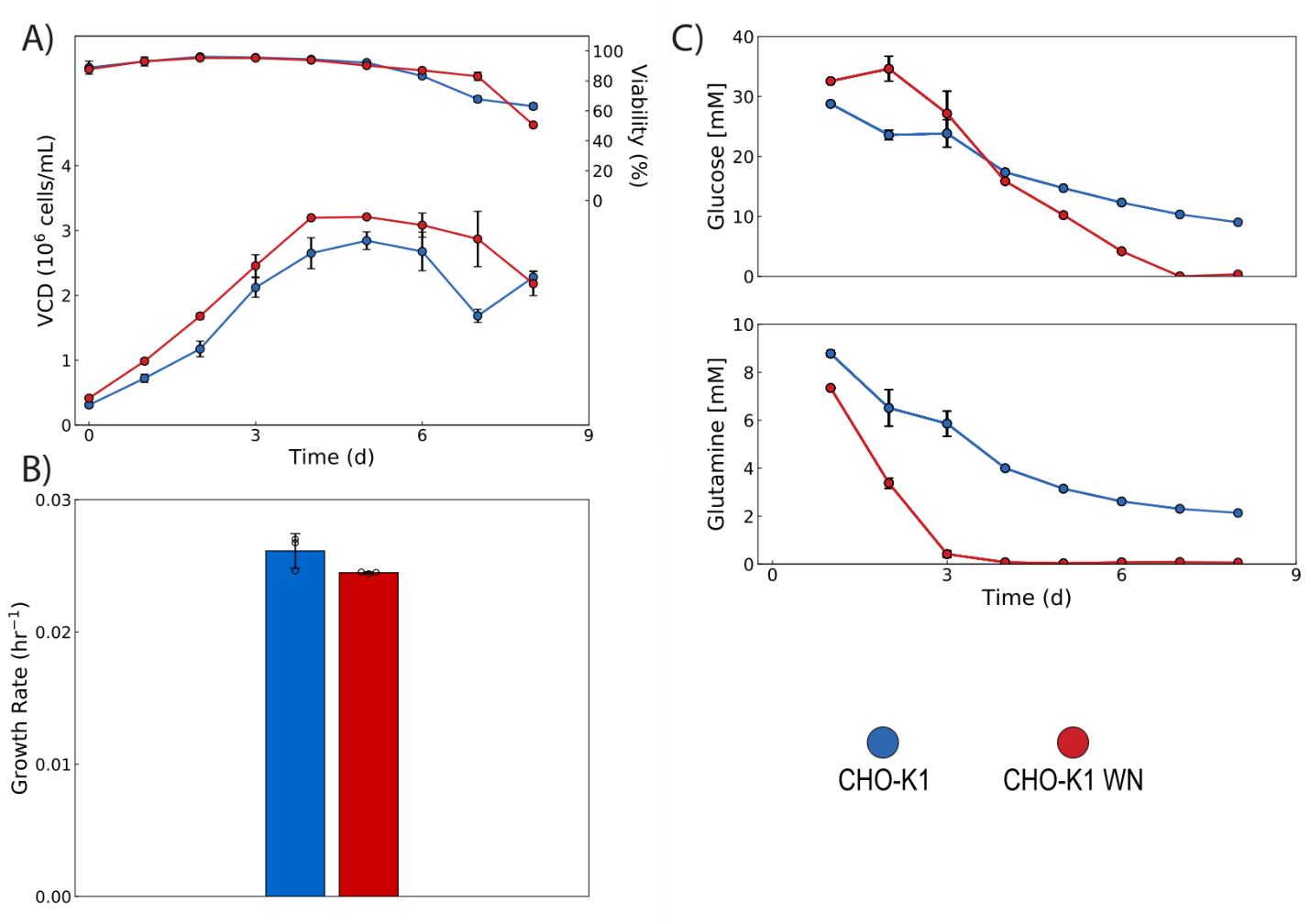

***Supplementary Figure 3: Warburg-null CHO-K1 batch culture***

*A Warburg-null CHO-K1 clone (CHO-K1 WN) and wildtype cells (CHO-K1) were grown in batch culture (n=3 shake flasks for each line). (A/B) Warburg-null cells grow as well and as quickly as wildtype cells. (C) CHO-K1 WN consumes glutamine more rapidly than the wildtype cell but consumes much less glucose while glutamine is available.*

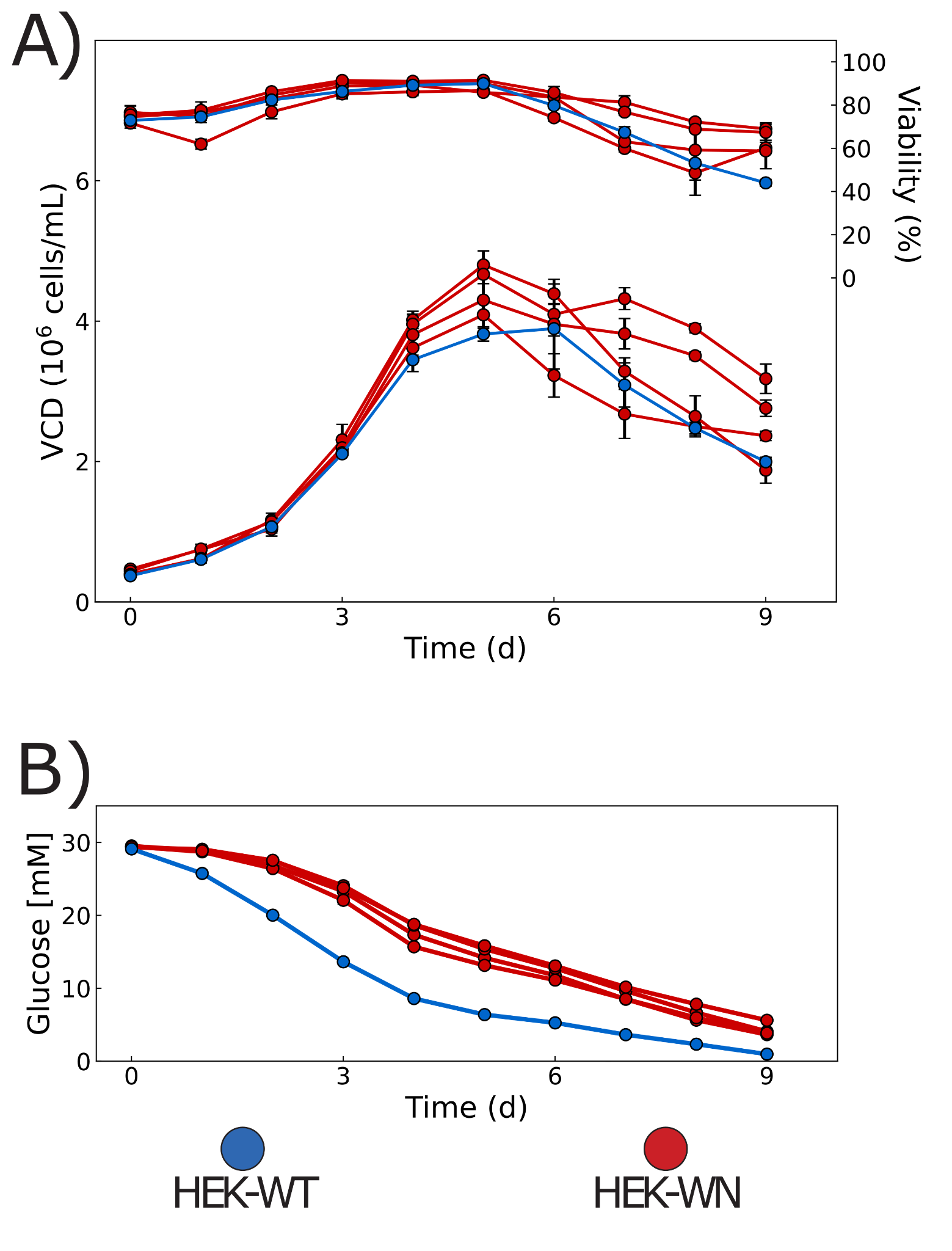

***Supplementary Figure 4: Warburg-null HEK 293F batch culture***

*(A) Warburg-null HEK 293F clones (HEK-WN) and wildtype (HEK-WT) were grown in batch culture and showed equivalent growth. (B) Warburg-null cells exhibit markedly reduced glucose consumption. For all panels, n=3 shake flasks for each cell line.*

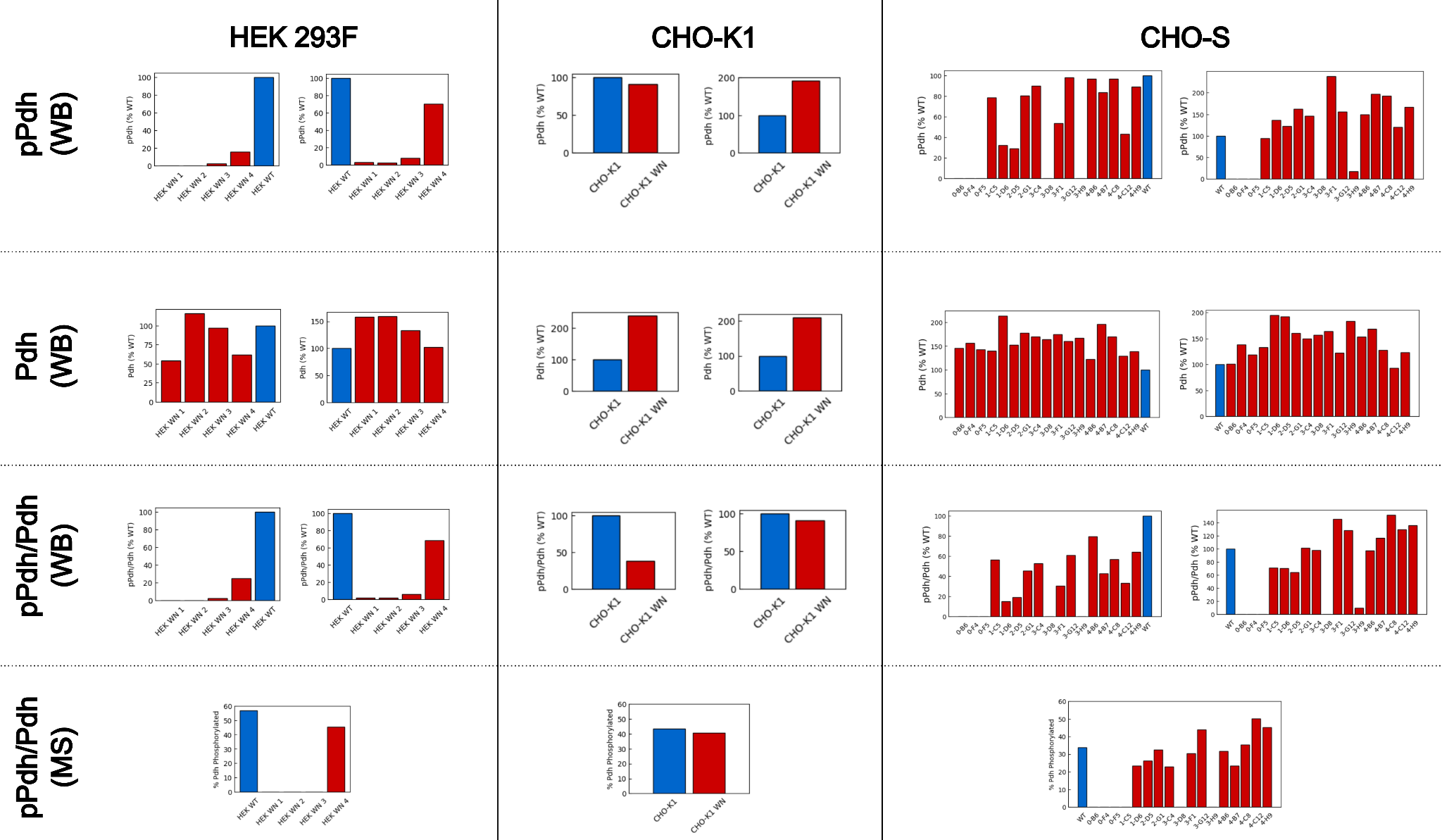

***Supplementary Figure 5: Quantification of pPdh and Pdh levels in Warburg-null clones***

*pPdh and Pdh levels were probed via Western blot (top two rows) in HEK 293F derived clones (left column), a CHO-K1 derived clone (middle column) and CHO-S derived clones (right column). The ratio of pPdh to Pdh from Western blot (3rd row, normalized to wildtype) and mass spectrometry (4th row, absolute quantification) is also shown. pPdh/Pdh from Western blots was calculated by taking the Vinculin-normalized intensity of the phosphorylated Pdh band divided by the Vinculin-normalized intensity of the Pdh band followed by normalization to the wildtype value. Western blots are depicted in Supplementary Figure 6. Mass spec values shown in Supplementary Table 10.*

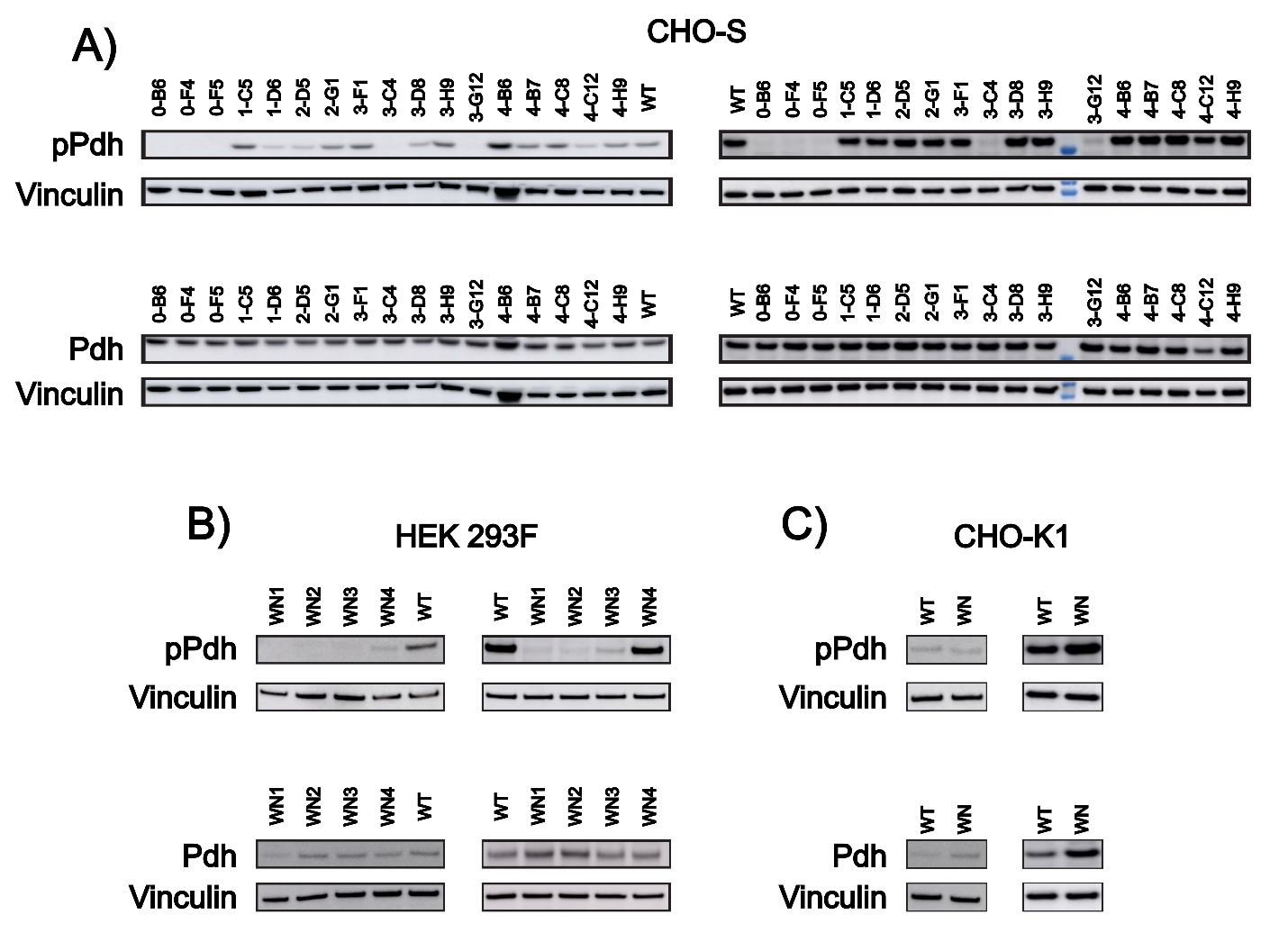

***Supplementary Figure 6: Western blots for pPdh and Pdh in Warburg-null clones***

*Duplicate blots for pPdh and Pdh in (A) CHO-S, (B) HEK 293F, and (C) CHO-K1 derived Warburg-null clones.*

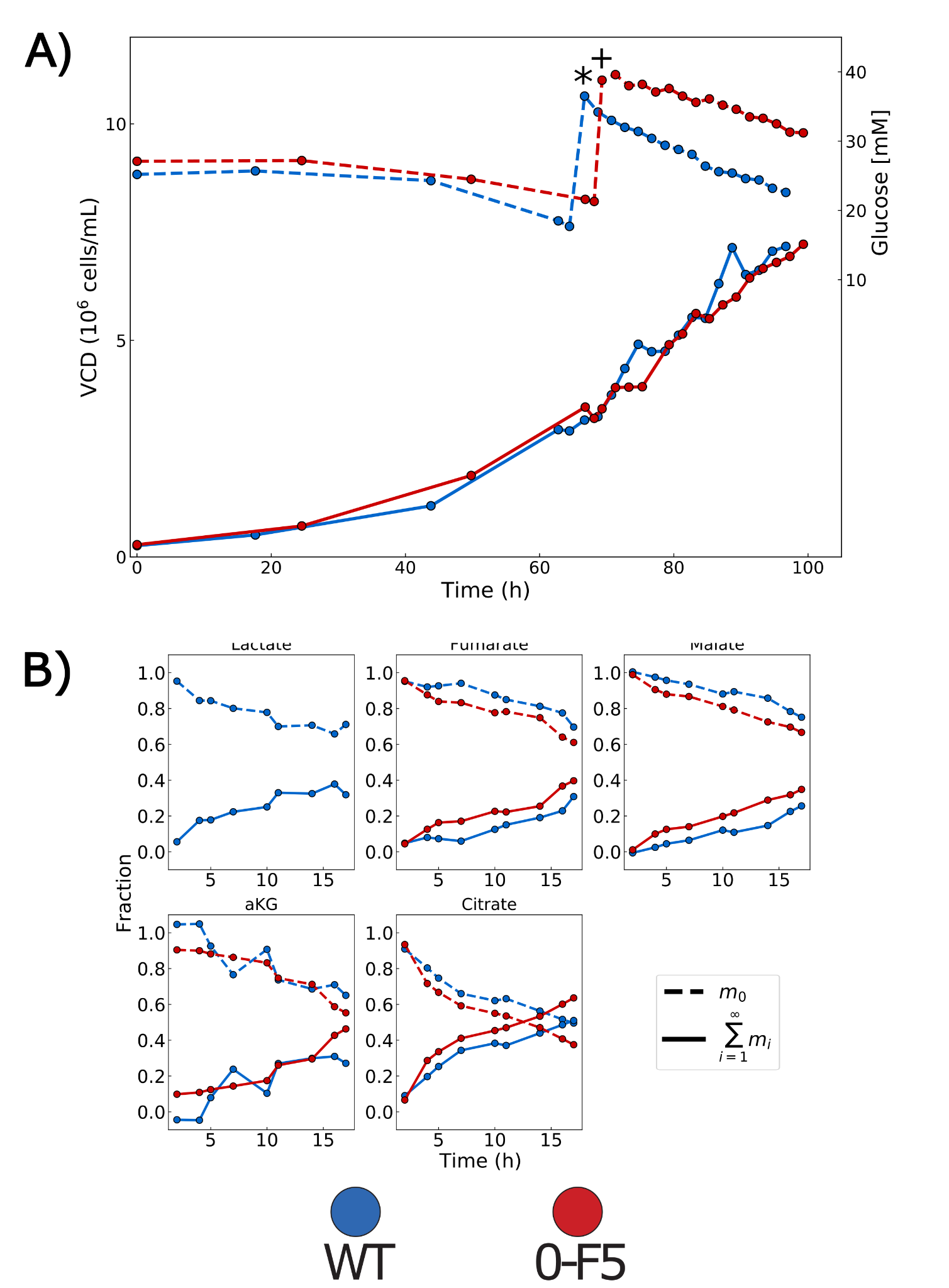

***Supplementary Figure 7: ^13^C labeling reveals increased TCA cycle activity in Warburg-null cells***

*(A) Cells were grown in bioreactors (n=1 bioreactor per cell line), when cells reached ~3x10^6^ cells/mL, ^13^C labeled glucose was spiked in and cultures were sampled every two hours starting at timepoints indicated by * and + for WT and 0-F5 cultures, respectively. (B) Unlabeled (m_0_) and sum of all labeled (*$\sum_{i=1}^{\infty} m_{i}$ *) metabolites are shown for relevant compounds. Time is calculated based on hours following ^13^C-labeled glucose addition.*
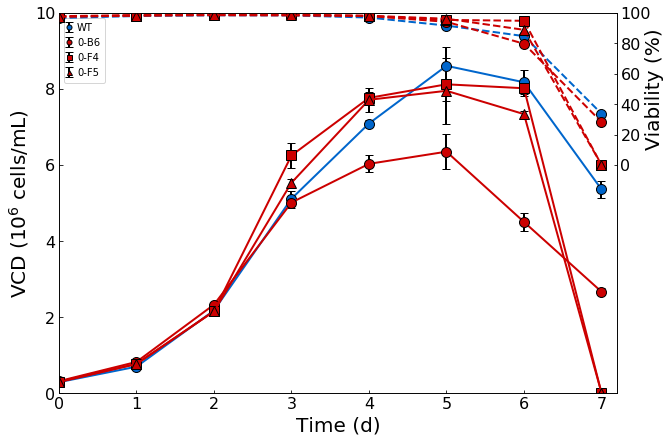

***Supplementary Figure 8: Growth profiles of Warburg-null CHO-S cell lines in shake flask batch culture***

*Clones 0-B6, 0-F4, 0-F5, and the wildtype CHO-S line were grown in shake flasks (n=3 shake flasks per cell line) for transcriptomics. All flasks were sampled on day 3 for RNA-Seq.*

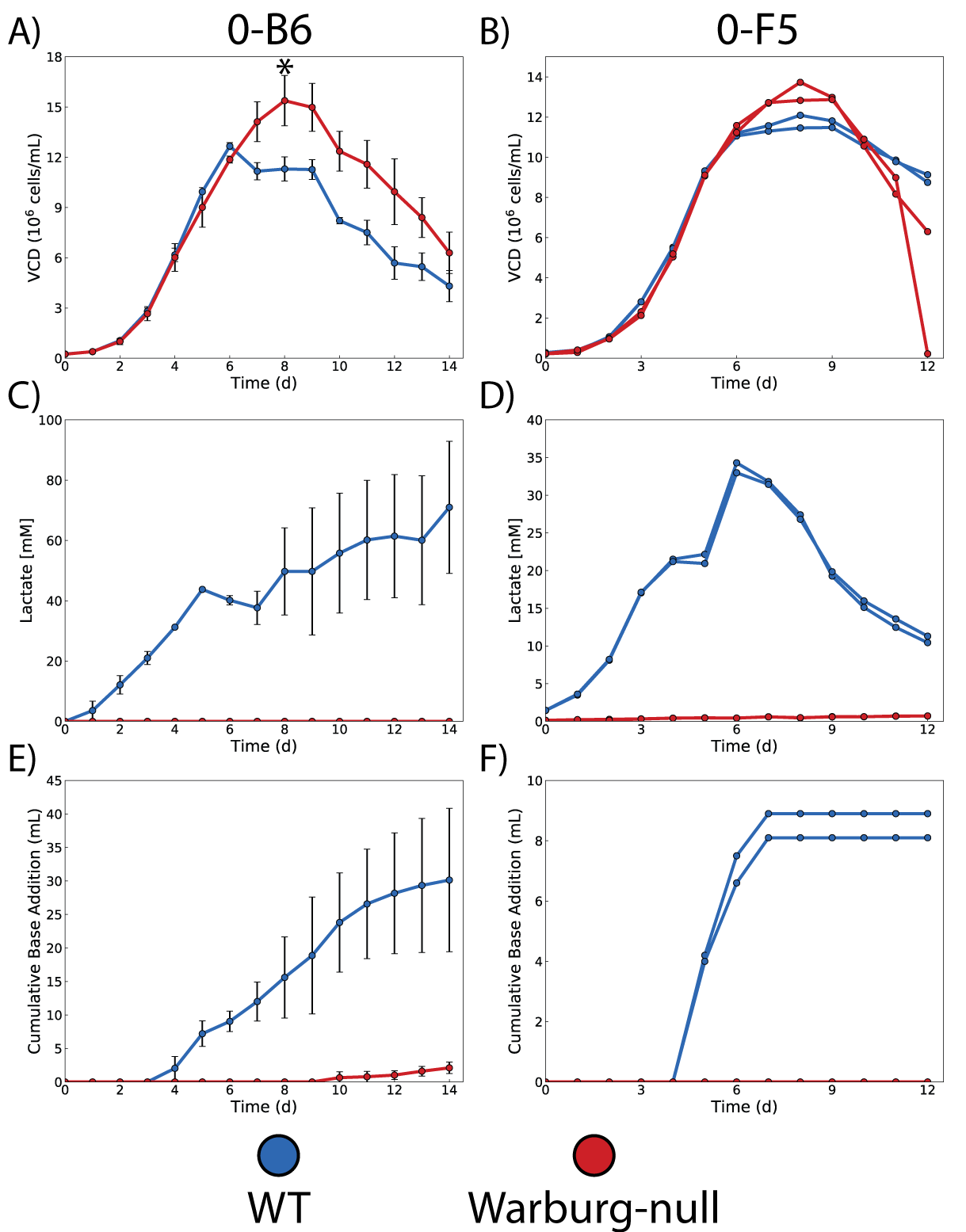

***Supplementary Figure 9: Elimination of the Warburg effect results in cells with a prolonged growth phase that do not require base addition to maintain culture pH***

*Two Warburg-null clones, 0-B6 (n=5 bioreactors, panels A, C, E) and 0-F5 (n=2 bioreactors, panels B, D, F), were grown in fed-batch culture with wildtype controls (n=3 bioreactors, panels A, C, E; n=2 bioreactors, panels B, D, F). (A/C) Knockout clones exhibited growth rate similar to WT but remain in exponential phase for a prolonged period of time (* indicates nutrient depletion for the 0-B6 culture, see Supplementary Figure 10). (B/D) Warburg-null clones remain non-lactogenic. (C/E) As a result of not secreting lactate, Warburg-null clones require less addition of base (1 M NaHCO_3_) over the course of culture to maintain pH at the appropriate levels (starting culture volume ~270 mL).*

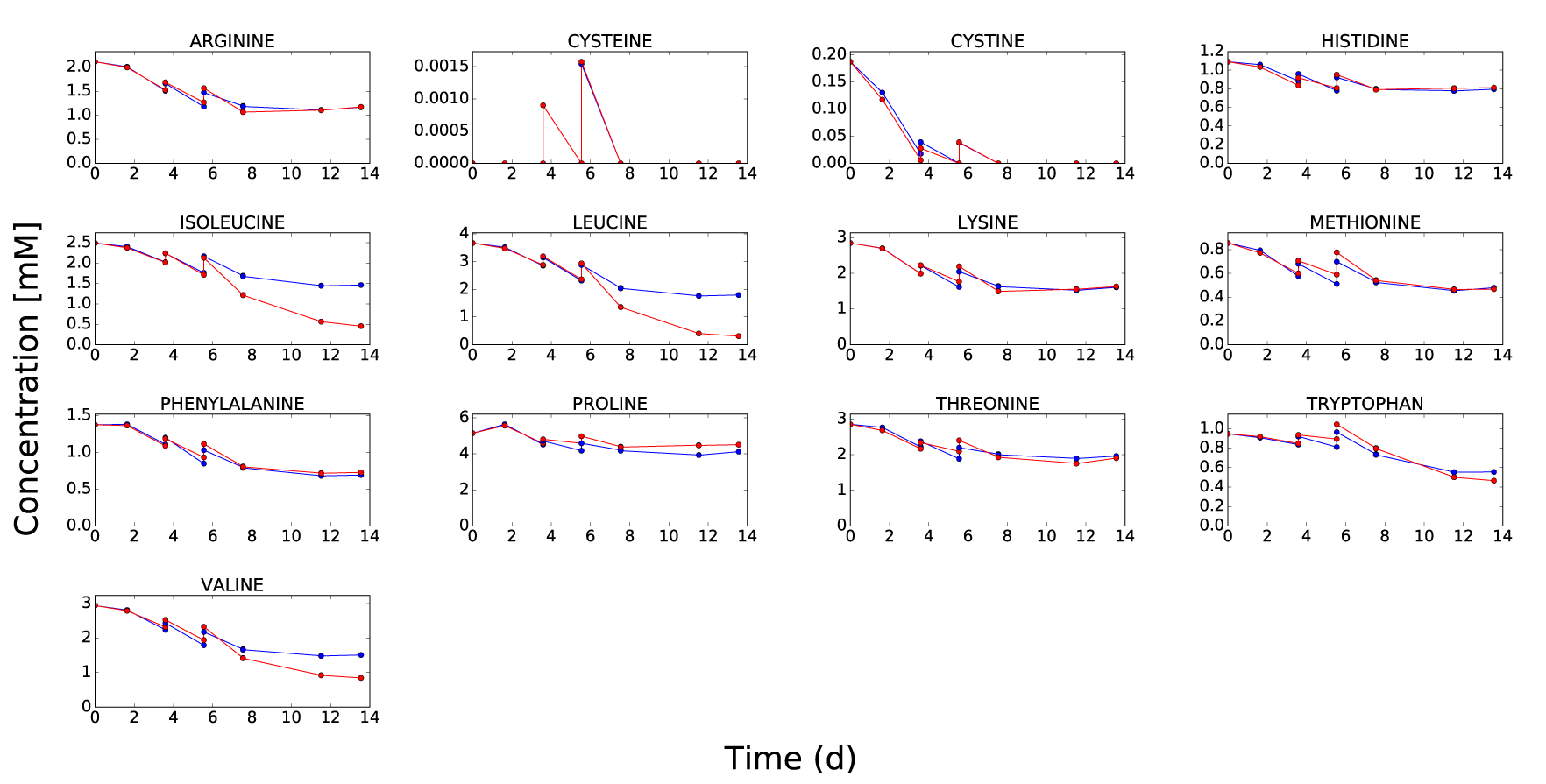

***Supplementary Figure 10: Time-course profiles of essential amino acids in fed-batch culture.***

*Amino acid concentrations from cultures shown in Supplementary Figure 9A (0-B6: red, n=5 bioreactors; WT: blue, n=3 bioreactors). Cysteine and cystine are depleted at day 8, at which point cells begin to die. Cells were fed on day 3, 4, 5, and 6 (see Supplementary Note for details on feeding strategy). Data shown as average concentrations before and after feed addition. Samples were analyzed via NMR by Spinnovation Biologics (Eurofins Spinnovation Analytical).*

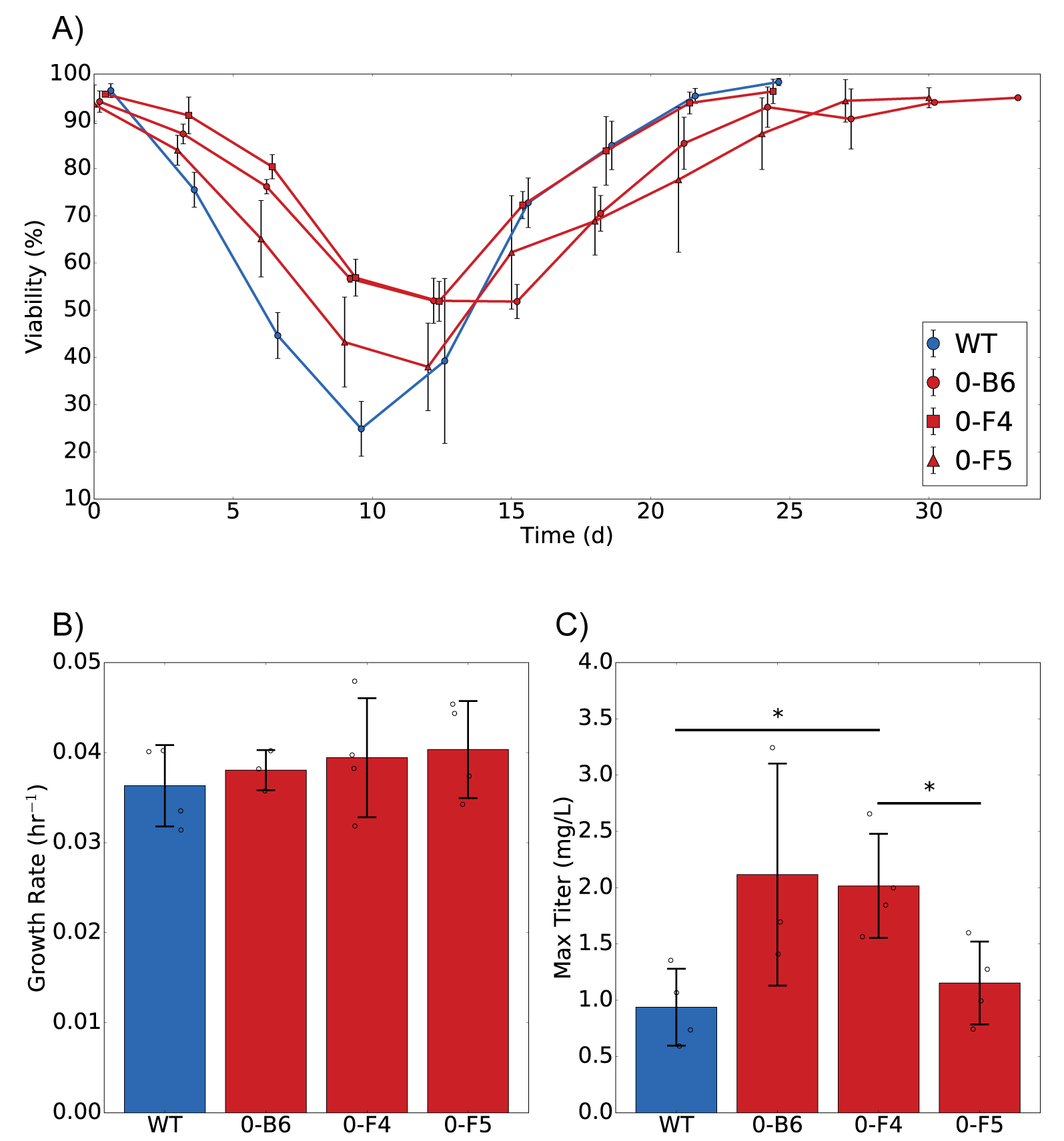

***Supplementary Figure 11: Eliminating the Warburg effect does not impact polyclonal pool generation****.*

*(A) Warburg-null clones show comparable recovery times to wildtype CHO-S cells when faced with a standard workflow for stable integration of a transgene. The resultant pools have indistinguishable growth (B) and protein production (C) characteristics. Thus, Ldha knockout clones are suitable for use as starting cell lines for biotherapeutic protein production. * indicates p<0.05 as determined by a two-sample two-tailed Welch’s t-test. Data shown as mean ± standard deviation. In all panels, n=3 shake flasks for 0-B6 and n=4 shake flasks for other lines (see Supplementary Figure 12). In (A), measurements taken every 3^rd^ day, slight offsets to displayed times for visual clarity.*

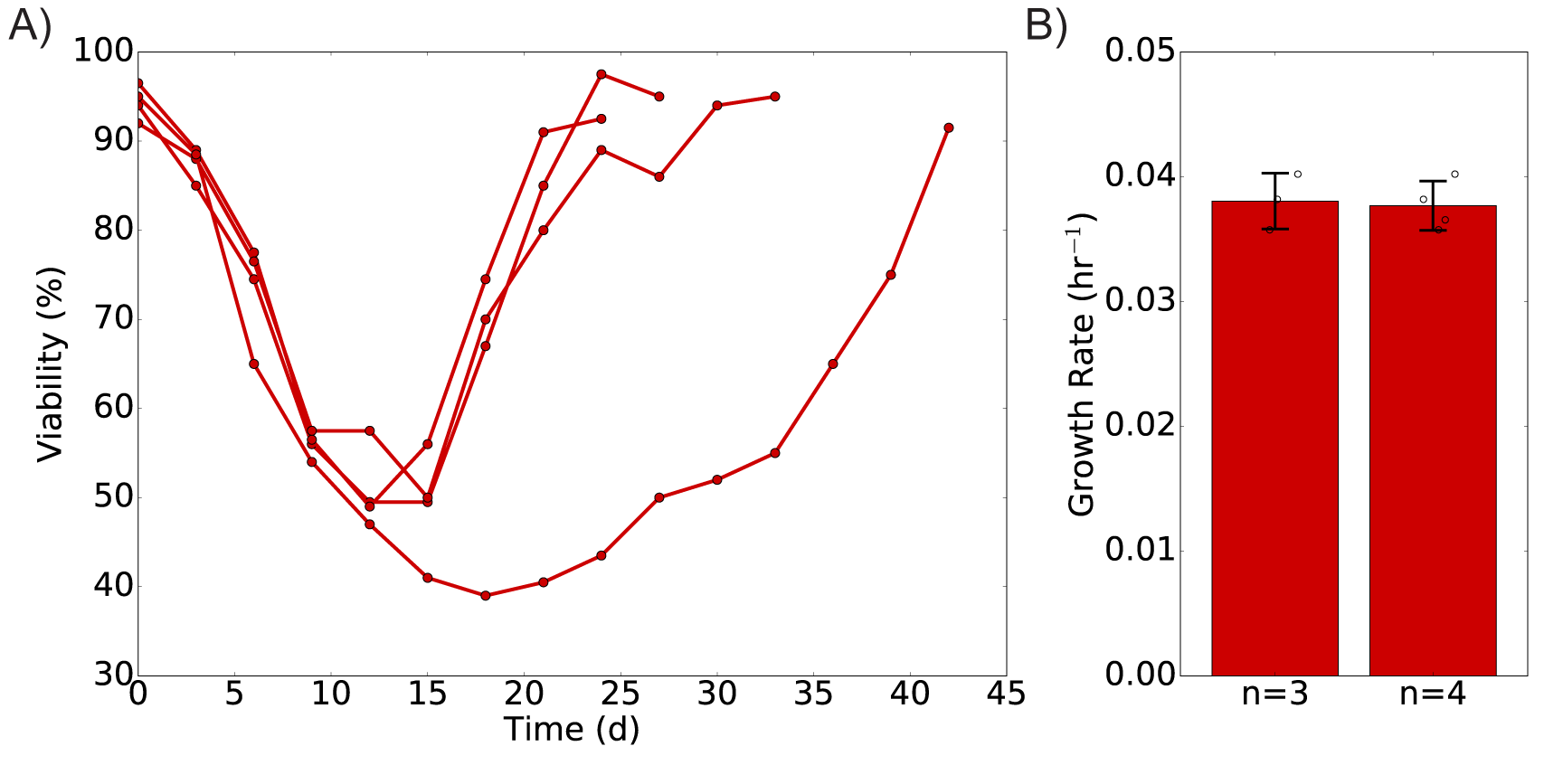

***Supplementary Figure 12: Outlier in pool generation process for clone 0-B6***

*During the cell line generation process (Supplementary Figure 11), one biological replicate of clone 0-B6 exhibited a drastically longer recovery time (A) for unknown reasons. While it is not included in the data presented in Supplementary Figure 1, preliminary characterization of the resultant pool with respect to growth (B) shows that it does not behave differently than any of the other 0-B6 derived clones (n=3 shake flasks, pools characterized in Supplementary Figure 11; n=4 shake flasks, pools characterized in Supplementary Figure 11 as well as the outlier from panel A).*

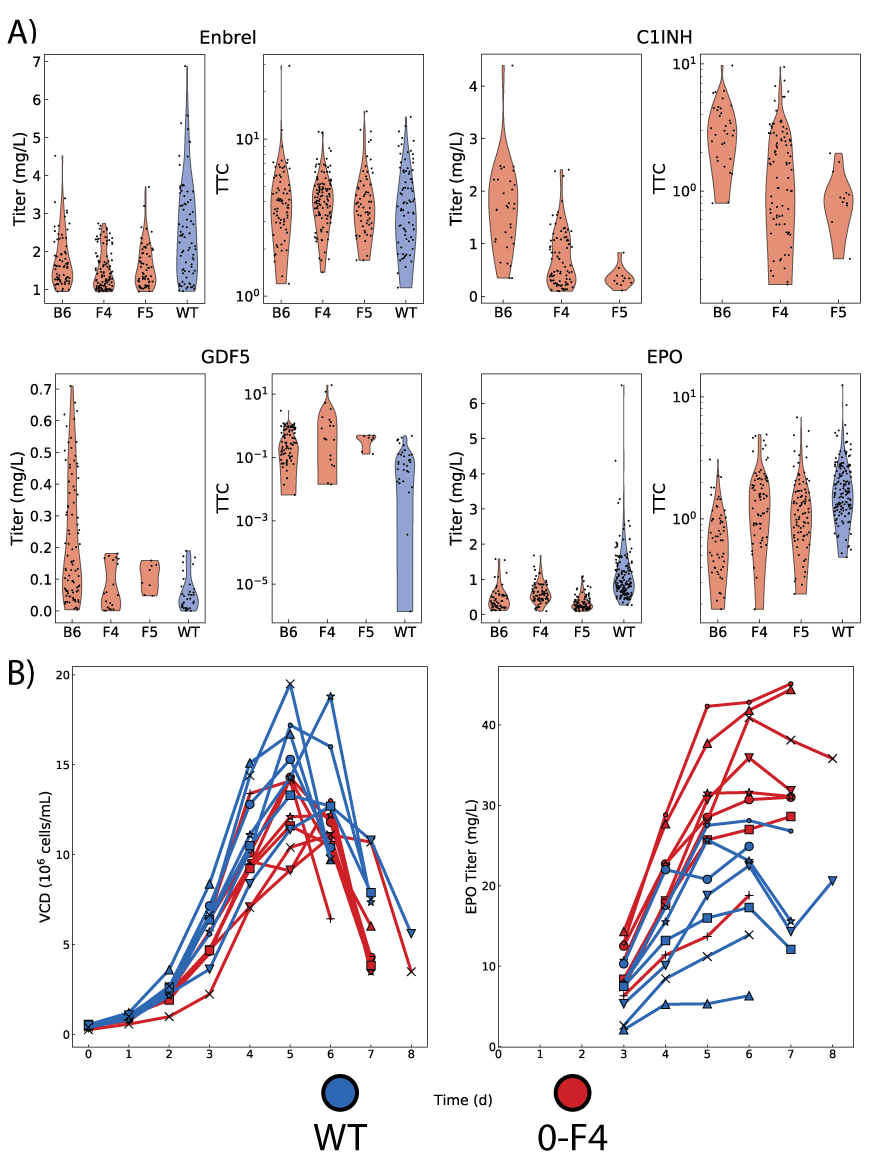

***Supplementary Figure 13: Warburg-null clones can be used for production of diverse biotherapeutic proteins***

*0-B6, 0-F4, 0-F5, and WT cells were subjected to a similar protocol as the one used in Supplementary Figure 11 to generate polyclonal pools producing erythropoietin (EPO), Enbrel, C1 esterase inhibitor (C1INH), and GDF5. Clonal protein-producing cells were isolated following FACS-assisted single cell sorting (using surface staining to enrich for high producers**^1^* *in the case of Enbrel). (A) All Warburg-null clones were able to successfully generate protein producing lines that behaved similarly to WT. Number of clones in (A)per starting cell line per protein are as follows: Enbrel-B6:72, F4:110, F5:63, WT:82; C1INH-B6:32, F4:86, F5: 12; GDF5-B6:88, F4:20, F5:8, WT:35; EPO-B6:55, F4:72, F5:87, WT: 158. The top 24 producing EPO clones derived from WT and 0-F4, as determined by the titer-to-confluence ratio (TTC)**^1^**, were expanded and reevaluated for production in suspension culture (12 well format). (B) Based on this, 8 Warburg-null clones and 7 WT derived clones were grown in batch culture (n=1 shake flask per cell line); again, clones derived from the Warburg-null cells produced as much—or more—protein than WT-derived clones.*

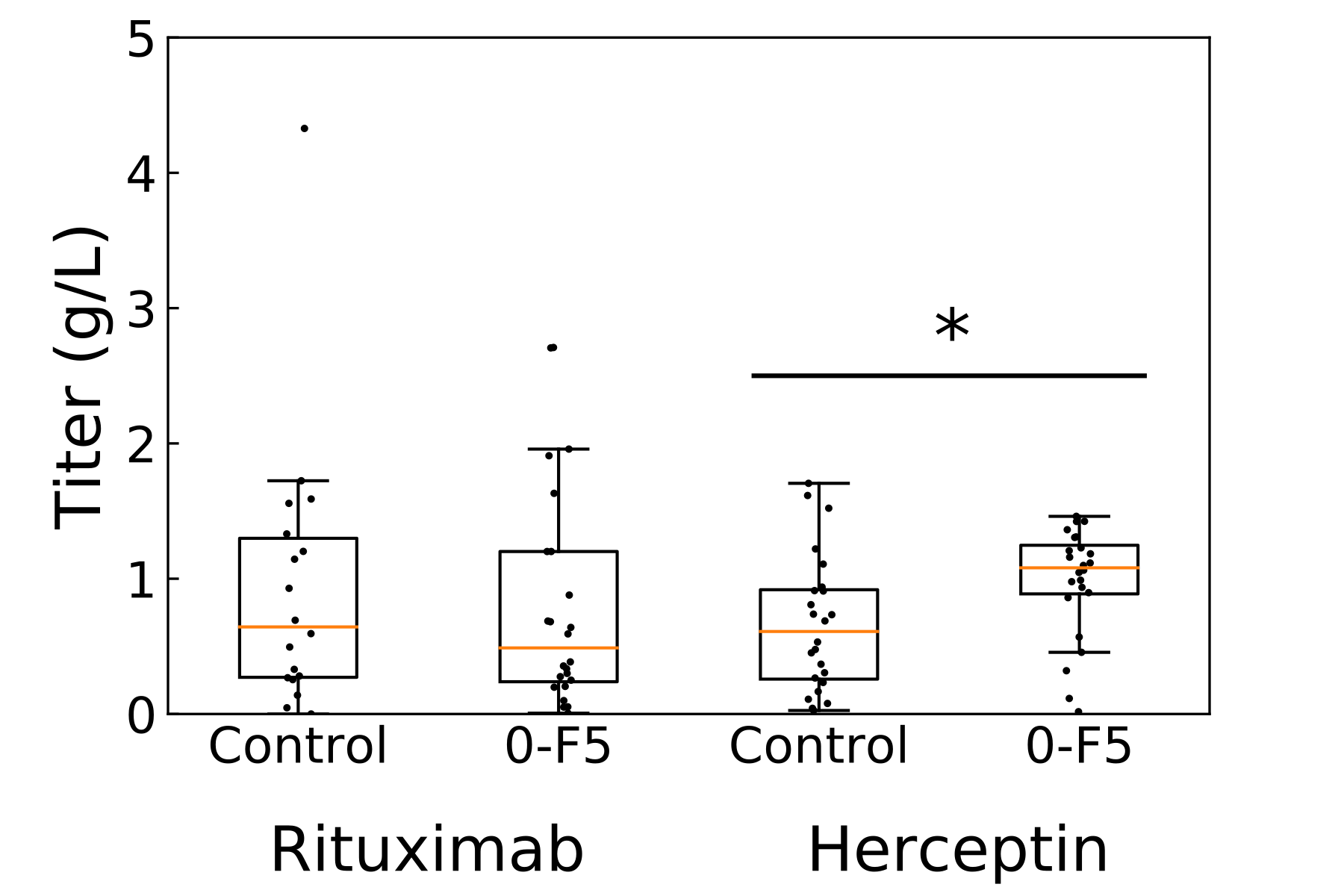

***Supplementary Figure 14: Fed-batch screening product titers***

*Day 10 titers in deepwell plates for clones generating Rituximab or Herceptin derived from a Warburg-null (0-F5) or Warburg-positive (Control) CHO cell line. N=24 clones for all samples except Control-Rituximab, where n=18. * indicates statistical significance (p<0.05) as determined by a Mann-Whitney U test.*

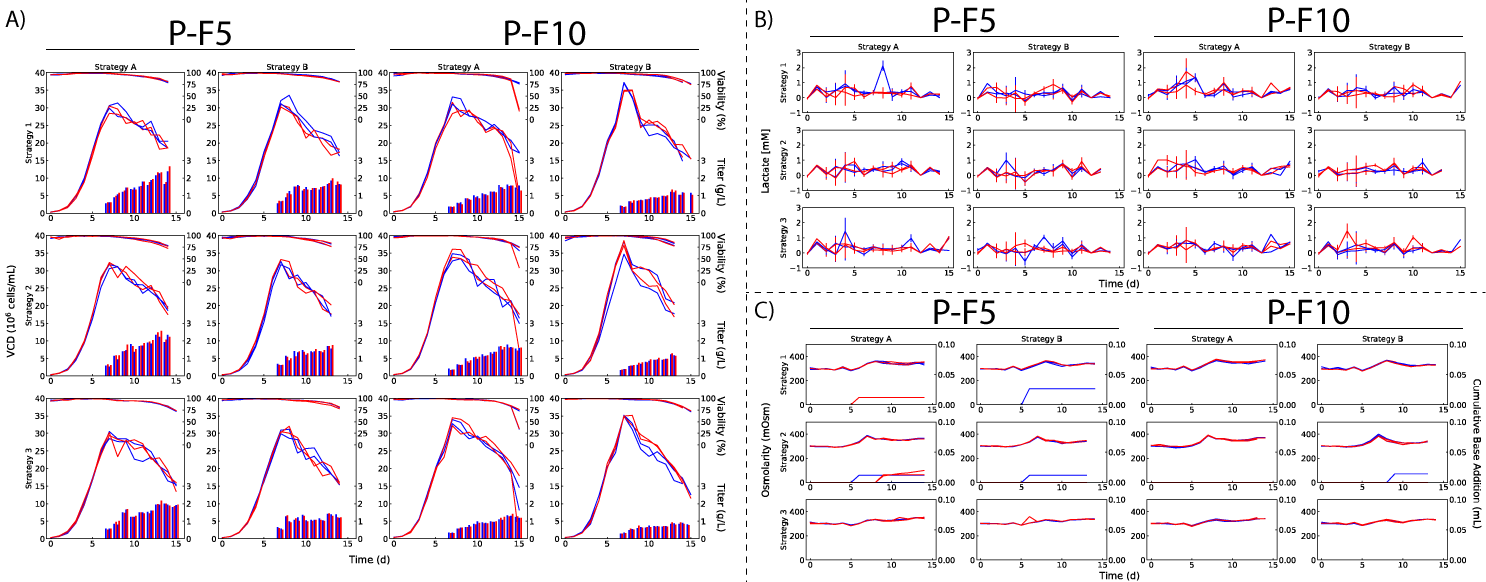

***Supplementary Figure 15: Warburg-null cells in ambr15 fed-batch***

*Two high-producing, Warburg-null Rituximab clones (Supplementary Figure 14), P-F5 and P-F10, were grown in ambr15 bioreactors with different feeding strategies and dissolved oxygen setpoints (blue-40%, red-50%; see Supplementary Note for details of feeding strategies; each unique combination was evaluated in duplicate bioreactors). (A) Cell density, viability, and product titer. (B) Lactate. (C) Osmolarity and base addition.*

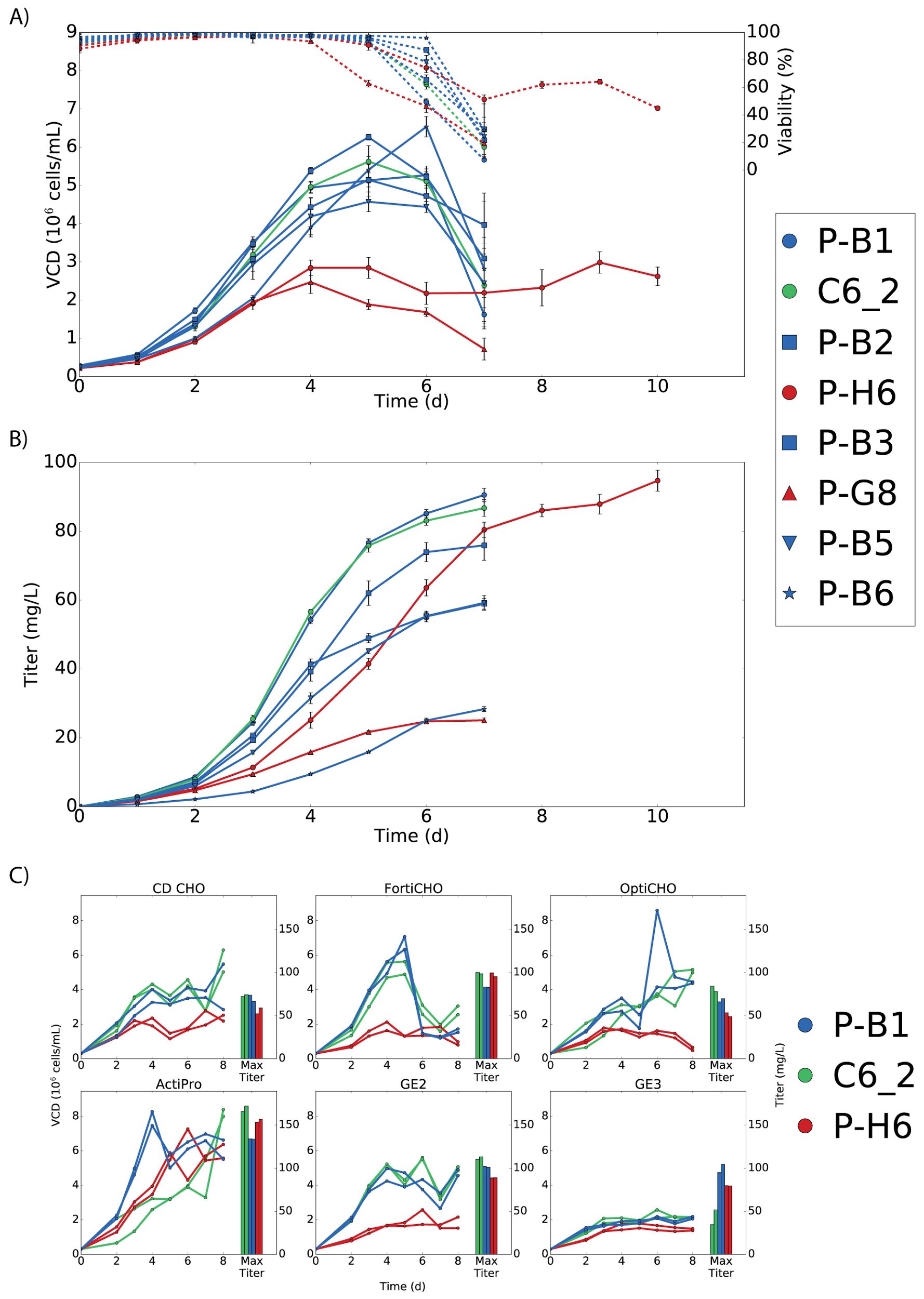

***Supplementary Figure 16: Batch-culture characterization and media optimization of clones derived from rituximab-producing line C6_2, following single-cell sorting.***

*Following single cell sorting and identification of Warburg-null clones, we grew the parental line, Warburg-null clones, and control clones in batch culture for preliminary characterization of the impact on growth and protein production. Significant variation is observed in both (A) growth profile and (B) rituximab titer both between engineered strains (i.e., KO vs. Mock vs. Parental) and within strains (e.g., P-B1 vs. P-B6) when grown in batch culture in CD CHO media (n=3 shake flasks for all lines). Data shown as mean ± standard deviation. Apparent poor growth of Warburg-null clones (P-H6/P-G8) was due to cell deposition on the flask surface at the liquid/air interface, artificially depressing cell counts). Since they had the highest product titer, we selected P-H6 and P-B1 for further characterization in fed-batch alongside the parental C6_2 line following a media screen to identify optimal media for growth and protein production (C). Media screening was carried out following the same protocol varying only the media (n=2 shake flasks for all lines). We tested CD CHO, FortiCHO (Gibco Cat. # A1148301), OptiCHO (Gibco Cat. # 12681011), ActiPro (GE Healthcare Cat. # SH31039.02), and 2 GE formulations, labeled here as GE2 (GE Healthcare Cat. # SH30871.02) and GE3 (GE Healthcare Cat. # SH30557.02). All media were supplemented with 1% antibiotic-antimycotic and 1 mL/L anti-clumping agent; all media except GE2 (which contained glutamine) was supplemented with 8 mM L-glutamine. Media screening revealed ActiPro permitted the best growth and protein production for all lines tested.*

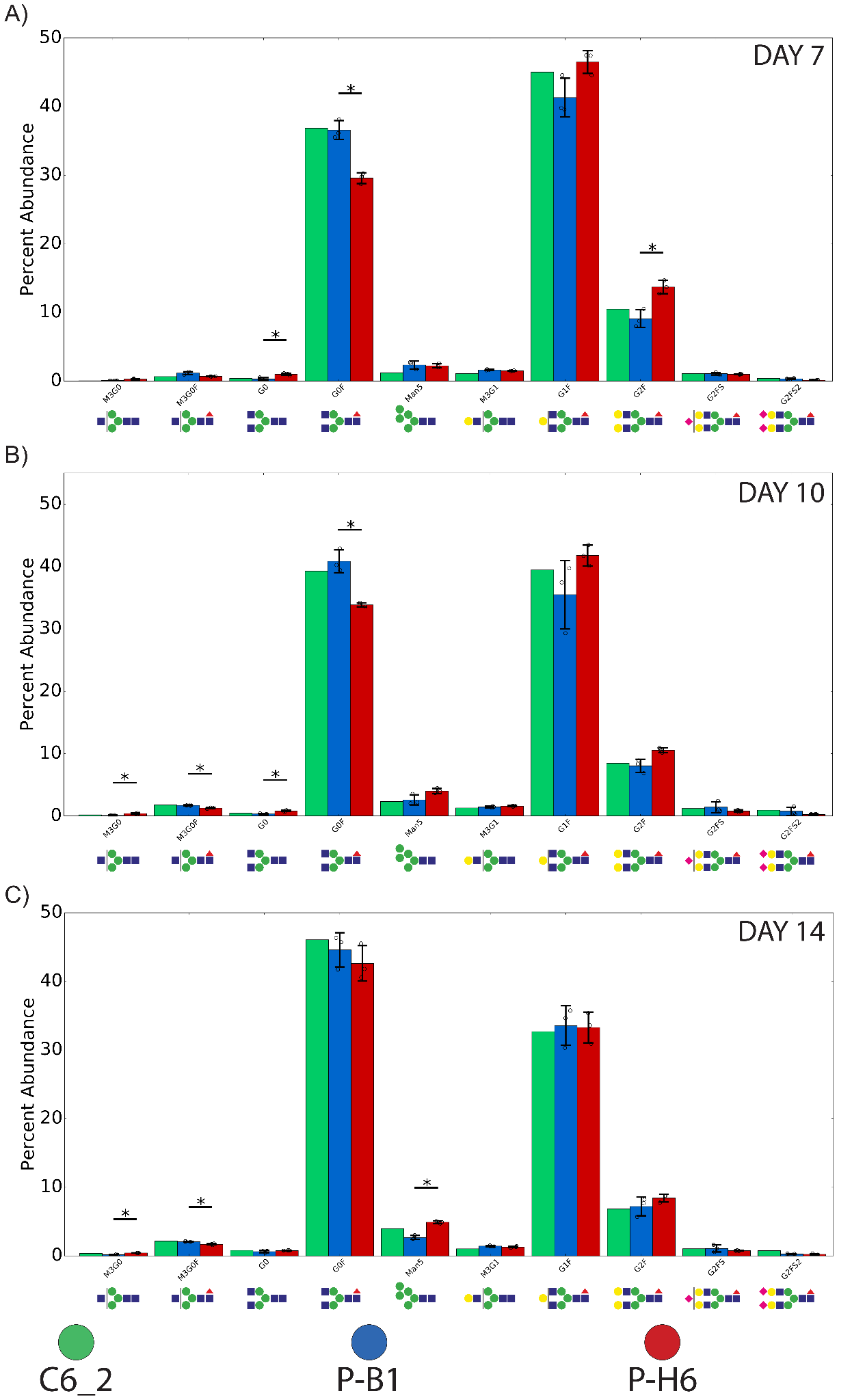

***Supplementary Figure 17: Rituximab glycoprofiles during fed-batch culture of parental and engineered cell lines.***

*Parental (C6_2, green, n=1 bioreactors), mock control (P-B1, blue, n=3 bioreactors), and Warburg-null (P-H6, red, n=3 bioreactors) cells producing rituximab in fed-batch (Figure 5D) had product glycoprofiles analyzed on day 7 (A), 10 (B), and 14 (C) of culture. * Indicates significance at an FDR of 0.05 as determined by the Benjamini–Hochberg method, with individual comparisons made using a two-sample two-tailed Welch’s t-test for mock vs. Warburg-null comparisons only. Data shown as mean ± standard deviation.*

|  | **Forward gRNA** | **Reverse gRNA** | **gRNA Sequence** | **PAM** | **Forward Primer** | **Reverse Primer** | **Protein Length** | **Cut Site (aa)** |
| --- | --- | --- | --- | --- | --- | --- | --- | --- |
| **Ldha** | GGAAAGGACGAAACACCGCTGGGCACTGATACCGACAGTTTTAGAGCTAGAAAT | CTAAAACTGTCGGTATCAGTGCCCAGCGGTGTTTCGTCCTTTCCACAAGATAT | GCTGGGCACTGATACCGACA | AGG | TCGTCGGCAGCGTCAGATGTGTATAAGAGACAGGTGCTCTCCTGTGGAAACATTG | GTCTCGTGGGCTCGGAGATGTGTATAAGAGACAGAGTTTCCATGCTGCCAATCACG | 332 | 216 |
| **Pdk1** | GGAAAGGACGAAACACCACTTCGACTACATCGCAGTTGTTTTAGAGCTAGAAAT | CTAAAACAACTGCGATGTAGTCGAAGTGGTGTTTCGTCCTTTCCACAAGATAT | ACTTCGACTACATCGCAGTT | TGG | TCGTCGGCAGCGTCAGATGTGTATAAGAGACAGGGGGTGTTAGGTAAATGCGC | GTCTCGTGGGCTCGGAGATGTGTATAAGAGACAGTTCTCTCATGACACAAGCAGT | 358 | 142 |
| **Pdk2** | GGAAAGGACGAAACACCGTCATTGTGCCGGTTCCGGAGTTTTAGAGCTAGAAAT | CTAAAACTCCGGAACCGGCACAATGACGGTGTTTCGTCCTTTCCACAAGATAT | GTCATTGTGCCGGTTCCGGA | TGG | TCGTCGGCAGCGTCAGATGTGTATAAGAGACAGGGGCTGGGGAGGATAGAGAA | GTCTCGTGGGCTCGGAGATGTGTATAAGAGACAGATGGAGATCCGGCTGAGGTA | 407 | 117 |
| **Pdk3** | GGAAAGGACGAAACACCATTGTGTTCGCCAGTCGTACGTTTTAGAGCTAGAAAT | CTAAAACGTACGACTGGCGAACACAATGGTGTTTCGTCCTTTCCACAAGATAT | ATTGTGTTCGCCAGTCGTAC | AGG | TCGTCGGCAGCGTCAGATGTGTATAAGAGACAGTGTTTCTCTGTGTTAATCTTAGGGA | GTCTCGTGGGCTCGGAGATGTGTATAAGAGACAGACTGAAGGGCGGTTCAACAA | 415 | 53 |
| **Pdk4** | GGAAAGGACGAAACACCTGGCGCTGCGCATCACGAAGGTTTTAGAGCTAGAAAT | CTAAAACCTTCGTGATGCGCAGCGCCAGGTGTTTCGTCCTTTCCACAAGATAT | TGGCGCTGCGCATCACGAAG | CGG | TCGTCGGCAGCGTCAGATGTGTATAAGAGACAGCCCAGGTCTCCTCTCACCAA | GTCTCGTGGGCTCGGAGATGTGTATAAGAGACAGGACCCTGAGTCCACAGCG | 412 | 3 |

***Supplementary Table 1: Oligos used for cloning gRNAs and primers for MiSeq analysis of CHO genes targeted in this study.***

*Sequences for generating gRNAs and primers used for engineering and amplicon sequencing. Pdk mutations introduce a frameshift upstream of the catalytic kinase domain, thus disrupting function**^2^**. Ldha mutations disrupt residues critical for substrate recognition and the Rossman fold domain necessary for NAD^+^/NADH binding**^3^**.*

| **Clone** | **Parent** | **Num. ID** | **Pdk1** | **Pdk2** | **Pdk3** | **Pdk4** | **Ldha** |
| --- | --- | --- | --- | --- | --- | --- | --- |
| 0-B6 | WT | 1 | X | X | X | X | X |
| 0-F4 | WT | 2 | X | X | X | X | X |
| 0-F5 | WT | 3 | X | X | X | X | X |
| 1-C5 | WT | 4 | X | X |  | X | X |
| 1-D6 | WT | 5 |  | X |  | X | X |
| 2-D5 | WT | 6 | X |  | ½ | X | X |
| 2-G1 | WT | 7 | X |  |  | X | X |
| 3-C4 | WT | 8 |  | X |  |  | X |
| 3-D8 | WT | 9 |  | X | X | X | X |
| 3-F1 | WT | 10 |  | X |  | X | X |
| 3-G12 | WT | 11 |  | X | X | X | X |
| 3-H9 | WT | 12 |  | X |  | X | X |
| 4-B6 | WT | 13 |  | X |  |  | X |
| 4-B7 | WT | 14 | X | X |  | X | X |
| 4-C8 | WT | 15 | X | X |  | X | X |
| 4-C12 | WT | 16 | ½ | X |  | ½ | X |
| 4-H9 | WT | 17 | ½ | X |  |  | X |
| P-H6 | C6_2 | - |  | X | X | X | X |
| P-G8 | C6_2 | - |  | X |  | X | X |
| P-F5 | 0-F5 | - | X | X | X | X | X |
| P-F10 | 0-F5 | - | X | X | X | X | X |

***Supplementary Table 2: Genotypes of isolated CHO Warburg-null clones.***

*Black box indicates wildtype alleles for the gene, X a full knockout, ½ a partial knockout. Shading of the cell indicates what portion of the observed indels are in-frame, white: all indels frameshift, red: all indels in-frame, yellow: some indels frameshift, some in-frame.*

| **Clone** | **Pdk1** | **Pdk2** | **Pdk3** | **Pdk4** | **Ldha** |
| --- | --- | --- | --- | --- | --- |
| 0-B6 | 1 | 1 | -8/4 | -2 | -8/-1 |
| 0-F4 | 1 | 1 | 1 | 1/-1 | -1 |
| 0-F5 | 1 | 1 | 101 | 1 | -1 |
| 1-C5 | -20/1 | 1 | - | -49/-1/1 | -4/-1 |
| 1-D6 | - | 4 | - | -17/-5 | -4/1 |
| 2-D5 | 1 | - | -24 | 1 | -2/1 |
| 2-G1 | 1 | - | - | 1/4 | -10 |
| 3-C4 | - | -10/1 | -15 | - | -4/4 |
| 3-D8 | - | -16/-10 | -8 | -24/-18 | -11/-2 |
| 3-F1 | - | 1 | -39 | -22/-21/1 | 1 |
| 3-G12 | - | 1 | -1 | 1 | -31/-19 |
| 3-H9 | - | 1 | - | -21 | -2/2 |
| 4-B6 | - | 1 | - | - | -4 |
| 4-B7 | -1/1 | -8/1 | - | -30/-17 | -88/-1 |
| 4-C8 | 1/12 | -16/1 | - | -24/-12 | -7/-2 |
| 4-C12 | 1 | 1 | - | 19 | -4/-1 |
| 4-H9 | 1 | -11 | - | - | -22 |
| P-H6 | - | -2/-1/1 | 1 | -13/1 | -2/-1 |
| P-G8 | - | -3/1 | - | -25/-19/-2/8 | -73/-10/-1 |

***Supplementary Table 3: Detailed genotypes of isolated CHO Warburg-null clones***

*Indel sizes in targeted genes for all characterized clones.*

| **Cell Line** | **Description** | **Reference** |
| --- | --- | --- |
| S (n=40) | Wildtype, 40 samples taken between mid-late exponential growth (72-180 hours of cultivation) from 4 biological replicates | PMID: 27883890 |
| K1 (n=2) | Wildtype, single timepoint | PMID: 27883890 |
| K1 (n=2) | Wildtype, single timepoint | PMID: 28074024 |
| K1 (n=2) | Wildtype, exponential and stationary | PMID: 28298216 |
| DG44 (n=6) | Wildtype, exponential and stationary phases with and without MEM NEAA in media. Two samples following NaBu treatment (24hr and 48hr post-treatment) | PMID: 28298216 |
| DXB11 (n=2) | Wildtype and Mre11 knockdown cells | PMID: 25672394 |

***Supplementary Table 4: Source for gene expression data in Figure 1.***

| **Clone** | **Parent** | **PDK1** | **PDK2** | **PDK3** | **PDK4** | **LDHA** |
| --- | --- | --- | --- | --- | --- | --- |
| CHO-K1 WN | CHO-K1 | ½ | X |  | ½ | X |

***Supplementary Table 5: Genotypes of the isolated CHO-K1 clone.***

*Black box indicates wildtype alleles for the gene, X a full knockout, ½ a partial knockout. Shading of the cell indicates what portion of the observed indels are in-frame, white: all indels frameshift, red: all indels in-frame, yellow: some indels frameshift, some in-frame.*

| **Clone** | **Parent** | **PDK1** | **PDK2** | **PDK3** | **PDK4** | **LDHA** |
| --- | --- | --- | --- | --- | --- | --- |
| CHO-K1 WN | CHO-K1 | 1 | 1 | - | -14 | -4/-1 |

***Supplementary Table 6: Detailed genotype of the isolated CHO-K1 clone.***

*Indel sizes in targeted genes for the characterized clone.*

|  | **Forward gRNA** | **Reverse gRNA** | **gRNA Sequence** | **PAM** | **Forward Primer** | **Reverse Primer** |
| --- | --- | --- | --- | --- | --- | --- |
| **LDHA** | GGAAAGGACGAAACACCGTTGTTGGGGTTGGTGCTGTGTTTTAGAGCTAGAAAT | CTAAAACACAGCACCAACCCCAACAACGGTGTTTCGTCCTTTCCACAAGATAT | TTGTTGGGGTTGGTGCTGT | TGG | TCGTCGGCAGCGTCAGATGTGTATAAGAGACAGTGGTTCCAAGTCCAATATGGCA | GTCTCGTGGGCTCGGAGATGTGTATAAGAGACAGGGGGTCAAGGTATGGGCTTC |
| **LDHB** | GGAAAGGACGAAACACCGTTCCAATCACGCGGTGTTTGTTTTAGAGCTAGAAAT | CTAAAACAAACACCGCGTGATTGGAACGGTGTTTCGTCCTTTCCACAAGATAT | TTCCAATCACGCGGTGTTT | GGG | TCGTCGGCAGCGTCAGATGTGTATAAGAGACAGaccacttGAGTCGCCATGTT | GTCTCGTGGGCTCGGAGATGTGTATAAGAGACAGCACCCCAAGCTGCCTAACA |
| **PDK1** | GGAAAGGACGAAACACCGTTTGTACCAATTGAACGGAGTTTTAGAGCTAGAAAT | CTAAAACTCCGTTCAATTGGTACAAACGGTGTTTCGTCCTTTCCACAAGATAT | TTTGTACCAATTGAACGGA | TGG | TCGTCGGCAGCGTCAGATGTGTATAAGAGACAGTGTTTCTGCGGCAAGAGTTG | GTCTCGTGGGCTCGGAGATGTGTATAAGAGACAGCCTTGAATAAAGTCCACAACTCCA |
| **PDK2** | GGAAAGGACGAAACACCGTCGATGCTGCCGATGTGTTGTTTTAGAGCTAGAAAT | CTAAAACAACACATCGGCAGCATCGACGGTGTTTCGTCCTTTCCACAAGATAT | TCGATGCTGCCGATGTGTT | TGG | TCGTCGGCAGCGTCAGATGTGTATAAGAGACAGGAAAGAGGAGGAAAGCCCGG | GTCTCGTGGGCTCGGAGATGTGTATAAGAGACAGACTGAGGAACGAGTCACCCT |
| **PDK3** | GGAAAGGACGAAACACCGTACTTAACCGCCCTTCAGTGTTTTAGAGCTAGAAAT | CTAAAACACTGAAGGGCGGTTAAGTACGGTGTTTCGTCCTTTCCACAAGATAT | TACTTAACCGCCCTTCAGT | GGG | TCGTCGGCAGCGTCAGATGTGTATAAGAGACAGAGGGAGAGATAATGCATGTGAGA | GTCTCGTGGGCTCGGAGATGTGTATAAGAGACAGAGAATGGTCTCGCTGCCAAA |
| **PDK4** | GGAAAGGACGAAACACCGGAATGTTGGCGAGTCTCACGTTTTAGAGCTAGAAAT | CTAAAACGTGAGACTCGCCAACATTCCGGTGTTTCGTCCTTTCCACAAGATAT | GAATGTTGGCGAGTCTCAC | AGG | TCGTCGGCAGCGTCAGATGTGTATAAGAGACAGTGGTTCACCACTTACCAGCTT | GTCTCGTGGGCTCGGAGATGTGTATAAGAGACAGAGGTTCAGAAAATGCATGTGAAAGa |

***Supplementary Table 7: Oligos used for cloning gRNAs and primers for MiSeq analysis of HEK genes targeted in this study.***

*Sequences for generating gRNAs and primers used for engineering and amplicon sequencing.*

| **Clone** | **Parent** | **PDK1** | **PDK2** | **PDK3** | **PDK4** | **LDHA** | **LDHB** |
| --- | --- | --- | --- | --- | --- | --- | --- |
| HEK-IM A | HEK-WT | X |  | X | X | X | ½ |
| HEK-WN 1 | HEK-IM A | X | X | X | X | X | X |
| HEK-WN 2 | HEK-IM A | X | X | X | X | X | X |
| HEK-IM B | HEK-WT | X |  | X | X | X | ½ |
| HEK-WN 3 | HEK-IM B | X |  | X | X | X | X |
| HEK-WN 4 | HEK-IM B | X | X | X | X | X | X |

***Supplementary Table 8: Genotypes of isolated HEK clones.***

*Black box indicates wildtype alleles for the gene, X a full knockout, ½ a partial knockout. Shading of the cell indicates what portion of the observed indels are in-frame, white: all indels frameshift, red: all indels in-frame, yellow: some indels frameshift, some in-frame.*

| **Clone** | **PDK1** | **PDK2** | **PDK3** | **PDK4** | **LDHA** | **LDHB** |
| --- | --- | --- | --- | --- | --- | --- |
| HEK-IM A | 1 | - | -25/-11/1 | -13/-2 | -6 | 1 |
| HEK-WN 1 | 1 | 1 | -25/-11/1 | -13/-2 | -6 | -24/-5 |
| HEK-WN 2 | 1 | -11 | -25/-11/1 | -13/-2 | -6 | -19/1 |
| HEK-IM B | -7/1 | - | -13/-8/1/2 | -5/-1/1 | -9/-6/1 | 1 |
| HEK-WN 3 | -7/1 | - | -13/-8/1/2 | -5/-1/1 | -9/-6/1 | -1/1/7 |
| HEK-WN 4 | -7/1 | 3/1 | -13/-8/1/2 | -5/-1/1 | -9/-6/1 | -2/1/3 |

***Supplementary Table 9: Detailed genotypes of isolated HEK clones.***

*Indel sizes in targeted genes for all characterized clones.*

| **Cell Line** | **Peptide** | | | **Total Protein Intensity** |
| --- | --- | --- | --- | --- |
|  | *Unmodified* | *Oxidized* | *Phosphorylated* |  |
| CHO-S WT | 42785.14453 | 54097.19141 | 49529.78125 | 4063248.996 |
| 0-B6 | 110616.7109 | 112428.9375 | 0 | 3964179.353 |
| 0-F4 | 119083.8203 | 137123.4922 | 0 | 5098887.572 |
| 0-F5 | 136981.2813 | 152637.6641 | 0 | 7932001.412 |
| 1-C5 | 80752.64844 | 123322.9531 | 0 | 8029411.353 |
| 1D-6 | 150792.3281 | 162872.0078 | 95935.03906 | 11573448.61 |
| 2-D5 | 114690.1953 | 141647.7891 | 91272.0625 | 9286223.691 |
| 2-G1 | 74854.59375 | 119417.0313 | 93606.91406 | 10789622.91 |
| 3-C4 | 135176.4531 | 149047.7031 | 85205.57813 | 8814867.332 |
| 3-D8 | 115893.1953 | 148648.5898 | 0 | 7127982.988 |
| 3-F1 | 61695.01172 | 82437.00977 | 63123.59375 | 4872413.178 |
| 3-G12 | 75329.57813 | 101668.0977 | 139366.3594 | 8168605.572 |
| 3-H9 | 217056.8594 | 264990.2578 | 0 | 12687876.81 |
| 4-B6 | 123904.0938 | 181597.6797 | 141503.9375 | 12082056.74 |
| 4-B7 | 139414.2813 | 101193.793 | 73438.35938 | 6722077.092 |
| 4-C8 | 33985.54688 | 46364.34375 | 43818.38281 | 2963947.677 |
| 4-C12 | 0 | 22402.20703 | 22534.19531 | 1510777.277 |
| 4-H9 | 77470.72656 | 90933.69141 | 138637.1719 | 7624644.111 |
| CHO-K1 WT | 31085.23633 | 33841.78125 | 49969.8125 | 2486229.773 |
| CHO-K1 WN | 94952.0625 | 98685.91797 | 131954.8906 | 8841696.517 |
| HEK 293F WT | 48701.19141 | 90108.92578 | 182745.3906 | 12890828.52 |
| HEK WN 1 | 360884.9375 | 312881.875 | 0 | 17207906.75 |
| HEK WN 2 | 279569.4063 | 256847.2031 | 0 | 14681783.29 |
| HEK WN 3 | 197921.1406 | 196148.5156 | 0 | 8303039.041 |
| HEK WN 4 | 65449.41406 | 50139.75977 | 95615.05469 | 4364142.785 |

***Supplementary Table 10: Mass spec based quantification of Pdh phosphorylation***

*Lysates from all cell lines were enriched for Pdh and subjected to mass spec to quantify the level of Pdh phosphorylation. Unmodified, oxidized, and phosphorylated peptides derived from Pdh were detected; percent phosphorylation was calculated by dividing the phosphorylated peptide intensity by the sum of all peptide intensities. Total protein intensity of the enriched lysate is also shown. Mass spectrometry data files have been deposited in the MassIVE repository with the following identifier: MSV000095049.*

**Supplementary Note: Fed-batch culture**

*ambr15 study (Figure 5A-C/Supplementary Figure 15)*

Cells were grown in ambr15 bioreactors as described in the main text with variations in feeding strategy and dissolved oxygen setpoint. If not listed here, all vessels were treated as described in the main text.

Three different initial feeding strategies were used. Different volumes of Cell Boost 7a and 7b (GE Healthcare) were added starting day 3 as a percentage of culture vessel volume:

Strategy 1:

Day 3: 3% 7a, 0.3% 7b

Day 5: 4% 7a, 0.4% 7b

Day 6: 4% 7a, 0.4% 7b

Day 7: 4% 7a, 0.4% 7b

Strategy 2:

Day 3: 1.5% 7a, 0.15% 7b

Day 4: 3% 7a, 0.3% 7b

Day 5: 4.5% 7a, 0.45% 7b

Day 6: 6% 7a, 0.6% 7b

Strategy 3:

Day 3: 2% 7a, 0.2% 7b

Day 5: 3% 7a, 0.3% 7b

Day 6: 3% 7a, 0.3% 7b

Day 7: 3% 7a, 0.3% 7b

Cell Boost 7a was added as 3 separate boluses in all strategies.

After the initial feeding strategy, two followup feeding strategies were implemented:

Strategy A:

Starting two days after the initial feeding strategy was completed (and on all subsequent days) cells were fed 3% vessel volume 1x Efficient Feed C+ (split into 3 separate boluses).

Strategy B:

One day after the initial feeding strategy was completed, 1.5 mL medium was removed and replaced with fresh medium (ActiPro supplemented with 1% antibiotic-antimycotic and 2 mL/L anti-clumping agent). On every subsequent day, the total sampling volume was replaced with fresh medium.

Half the vessels were maintained at a DO setpoint of 40%, the other half at 50%. Every combination of DO setpoint, initial feeding strategy, and followup feeding strategy was run in duplicate.

*Nonproducer study (Supplementary Figure 9)*

Cells were grown from a seed density of 2.5x10^5^ cells/mL in DASGIP bioreactors with a starting volume of 270 mL at a temperature of 37°C, agitated at 200 rpm using pitched blade impellers. The pH was maintained continuously at 7.10±0.02 with 1 M sodium bicarbonate or CO_2_. Dissolved oxygen was maintained at 40% using pure oxygen, air, or a mixture, as needed. A 2222 mM glucose solution and a 200 mM glutamine solution were used to control glucose and glutamine levels at the desired levels via a once-daily feeding based on the measured concentration, doubling time, and calculated consumption rates. Complex feed addition was started when cell concentration was approximately 1.4-2.2x10^6^ cells/mL and continued for 4 days, feeding increasing volumes each successive day. Cultures were sampled daily for cell growth and viability using the NucleoCounter NC-200 Cell Counter and metabolite concentrations using the BioProfile 400. Process variations for the different experiments are detailed below:

*Clone 0-B6*

Cells were maintained in CD CHO medium supplemented with 8 mM L-glutamine, 1% antibiotic-antimycotic (Life Technologies Cat. # 15240062), and 1 mL/L anti-clumping agent. 0.04% Pluronic F-68 (Life Technologies Cat. # 24040032) was added to each reactor prior to inoculation and supplemented as needed over the course of culture. Glucose was maintained between approximately 24-40 mM while additional glutamine was not supplemented over the course of culture. Efficient Feed B (Thermo Fisher Cat. # A1024001) was added at the following days/volumes:

Day 3: 4 mL

Day 4: 8 mL

Day 5: 12 mL

Day 6: 16 mL

*Clone 0-F5*

Cells were maintained in CD CHO medium supplemented with 8 mM L-glutamine, 1% antibiotic-antimycotic, and 1 mL/L anti-clumping agent. Antifoam C was added as needed over the course of culture. Glucose was maintained between 10-24 mM. Glutamine was maintained between 1-3 mM. Efficient Feed B was added at the following days/volumes:

Day 3: 4 mL

Day 4: 8 mL

Day 5: 12 mL

Day 6: 16 mL
